# Kingdom-Wide CRISPR Guide Design with ALLEGRO

**DOI:** 10.1101/2025.02.13.638206

**Authors:** Amirsadra Mohseni, Reyhane Ghorbani Nia, Aida Tafrishi, Xin-Zhan Liu, Jason E. Stajich, Ian Wheeldon, Stefano Lonardi

## Abstract

Designing CRISPR single guide RNA (sgRNA) libraries targeting entire kingdoms of life will significantly advance genetic research in diverse and underexplored taxa. Current sgRNA design tools are often species-specific and fail to scale to large, phylogenetically diverse datasets, limiting their applicability to comparative genomics, evolutionary studies, and biotechnology. Here, we present ALLEGRO, a combinatorial optimization algorithm able to design minimal, yet highly effective sgRNA libraries targeting thousands of species. Leveraging integer linear programming, ALLEGRO identified compact sgRNA sets simultaneously targeting several genes of interest for over 2,000 species across the fungal kingdom. We experimentally validated the sgRNAs designed by ALLEGRO in *Kluyveromyces marxianus, Komagataella phaffii*, and *Yarrowia lipolytica*. In addition, we adopted a generalized Cas9-Ribonucleoprotein delivery system coupled with protoplast transformation to extend ALLEGRO’s sgRNA libraries to other untested fungal genomes, such as *Rhodotorula araucariae*. Our experimental results, along with cross-validation, show that ALLEGRO enables efficient CRISPR genome editing, supporting the development of universal sgRNA libraries applicable to entire taxonomic groups.

## Introduction

Clustered regularly interspaced short palindromic repeats (CRISPR) and their associated proteins, known as CRISPR-Cas systems, are innate bacterial and archaeal defense mechanisms that use CRISPR RNAs (crRNAs) to detect and induce double-stranded breaks into foreign nucleic acids, consequently silencing them (1–3). Over the past decade, these defense mechanisms have been “hijacked” into genome editing systems to great success. However, most research has largely focused on model organisms, leaving a gap in accessibility for researchers working on diverse taxonomic groups and non-model organisms. Additionally, current single guide RNA (sgRNA or simply *guide*) design tools are often species-specific, with limited flexibility to address the challenges of cross-species variation in target and background sequences (4). These constraints hinder the development of scalable solutions for editing multiple genes across diverse genomes, particularly when addressing large-scale studies or species outside the conventional models. The ability to design effective CRISPR libraries across species is crucial for advancing genome editing in underexplored taxa. Expanding CRISPR applications to diverse species enables the integration of unique organisms into engineering and discovery pipelines, unlocking novel genetic and enzymatic pathways previously inaccessible with traditional approaches. However, designing and cloning individually tailored sgRNAs for multiple genes across numerous genomes remains experimentally burdensome and inefficient, making large-scale genomic screens tedious. This highlights the need for design tools that can generate a minimal set of guides capable of efficiently covering all desired targets across multiple species.

The problem of finding the smallest set of sgRNAs that targets a given set of genes in a given set of species is a type of the *Set Covering Problem*. In this combinatorial optimization problem, given a collection of elements (e.g., genomes or genes) and a collection of sets containing the elements (e.g., sgRNAs), the goal is to select the smallest number of sets whose union covers all the elements. This problem has been extensively studied in Computer Science, and is known to be NP-hard, which is a class of problems that includes the most computationally demanding problems to solve optimally (5). Designing a compact guide library across many genomes has been previously studied and a few tools have been designed for this purpose. For instance, Endo et al. leveraged the mismatch tolerance of Cas9 guides in the PAM-distal region to design an sgRNA capable of targeting homologous genes of interest in the rice genome (6). CRISPR MultiTargeter similarly identifies common guides using multiple sequence alignments as input, though it does not optimize for the smallest guide set (7). CRISPys is another tool designed for creating guide libraries using various strategies, including one functionality that identifies minimal sets of guides targeting small sized gene sets (8). The tool that best addresses the guide design challenge is MINORg (9), which can identify the minimal set of guides targeting each input sequence but only for a limited size set of genes and guides. While the algorithms employed by the available methods can be effective in designing a minimal guide library for relatively small datasets, they prove to be computationally inefficient and are impractical for large-scale data. To facilitate the design of guide libraries that cover a large number of diverse species, here we present ALLEGRO (*Algorithm for a Linear program Enabling Guide RNA Optimization*), a time- and memory-efficient algorithm capable of designing a minimal guide library for thousands of organisms within minutes. ALLEGRO uses advanced combinatorial optimization techniques to formulate and solve a modified version of the minimum set covering problem using integer linear programming. Designed for flexibility and ease of use, ALLEGRO allows users to (i) select between two sgRNA design strategies, called *tracks*, (ii) filter guides based on sequence features, and (iii) cluster sgRNAs to further reduce the size of the libraries. These features, among others, enable the creation of guide libraries applicable across entire biological kingdoms.

To validate the sgRNA libraries designed by ALLEGRO, we conducted a series of CRISPR-Cas9 knockout experiments across various fungal species, including *Kluyveromyces marxianus, Komagataella phaffii, Yarrowia lipolytica*, and *Rhodotorula araucariae*. The tested sgRNAs were designed to target a set of auxotrophic genes to enable screening. Our validation experiments confirmed that ALLEGRO designs highly efficient and precise sgRNAs, yielding robust gene editing outcomes in all tested species. In addition, we developed a generalized approach for applying ALLEGRO sgRNA libraries to other fungal genomes, leveraging a general Cas9 delivery method based on Cas9-Ribonucleoprotein (RNP) and protoplast transformation. These results demonstrate ALLEGRO’s capability to design effective and functional sgRNAs for genome engineering across diverse fungal species, facilitating broader applications in microbiology and synthetic biology.

## Materials and Methods

### Computational Composition

The input to ALLEGRO is a set of species and a set of genes of interest. In this work, we apply ALLEGRO to the fungal kingdom and a several genes of interest, but the algorithm is flexible and can be adopted for other species and genes or genomes. First, ALLEGRO determines orthogroups (set of genes descended from a single gene in the last common ancestor of the species) across all input species for the genes of interest. The orthogroups are determined using reciprocal best hits with DIAMOND (10). ALLEGRO then computes predicted efficiency scores for all Cas9 guides in the orthogroups using the Ucrispr guide design algorithm (11). Finally, ALLEGRO determines the smallest library of guides that maximizes the cutting efficiency while targeting all genes in the set. An overview of the workflow is illustrated in Figure 1.

**Fig. 1.**
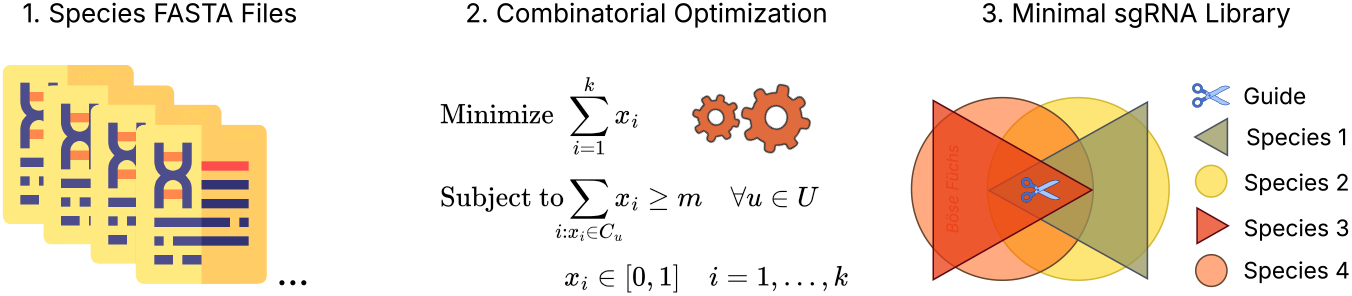
ALLEGRO’s workflow. **Step (1)** Given the gene sequence or the genome of hundreds to thousands of input species, ALLEGRO extracts Cas9 target sequences. **Step (2)** ALLEGRO builds and solves an (integer) linear program involving millions of variables. **Step (3)** The optimal solution of the linear program determines the sgRNA library with minimal size that covers all targets.

ALLEGRO leverages advanced techniques developed for combinatorial optimization to find the smallest set of sgRNAs that can target a set of genes of interest in a given set of species. The set of guides can be designed under user-defined constraints, which we call *tracks*. ALLEGRO allows users to choose two types of tracks, namely *A* for *any* and *E* for *each*. Tracks are also further characterized by the multiplicity factor *m*. When users choose track *A*_*m*_, ALLEGRO’s solution guarantees that any of the genes in every species is targeted at least *m* times by the guides in the library. When users choose track *E*_*m*_, ALLEGRO’s solution guarantees that each gene must be targeted at least *m* times in each species. A detailed description of the mathematical formulation, tracks, and multiplicity is provided in **Supplementary Notes**, and a more detailed overview is provided in **Supplementary Figure 1**.

### Experimental Validation

#### Microbial Strains and Culturing

All strains used in this study are listed in **Supplementary Table 1** *Yarrowia lipolytica* PO1f (*MatA, leu2-270, ura3-302, xpr2-322, axp-2*), *Kluyveromyces marxianus* CBS6556 (*ura3Δ*), *Komagataella phaffii* GS115 (*his4::CAS9*), and *Rhodotorula araucariae* NRRL Y-17376 were used in ALLEGRO guide design validation experiments. Unless stated otherwise, yeast cultures were grown in 14 mL polypropylene tubes or 250 mL baffled flasks at 30°C or 37°C with shaking at 225 rpm. Under non-selective conditions, yeast strains were cultivated in YPD medium (1% Bacto yeast extract, 2% Bacto peptone, 2% glucose).

**Table 1.**
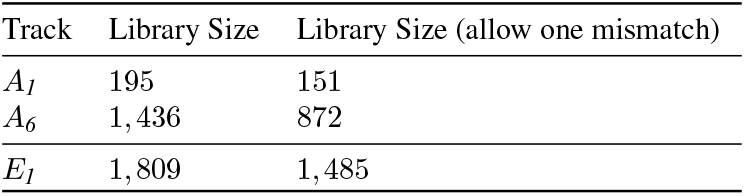
The size of ALLEGRO’s libraries that target six auxotrophic genes in 2,263 fungal species for tracks*A*_*1*_, *A*_*6*_, and *E*_*1*_. Allowing one mismatch in the PAM-distal region of the guides significantly reduces the library sizes. These experiments exclude sgRNA spanning intron-exon boundaries.

*Y. lipolytica* and *K. marxianus* transformants were initially propagated in synthetic defined media lacking uracil (SD-ura; 0.67% Difco yeast nitrogen base (YNB) without amino acids, 0.069% CSM-ura [Sunrise Science, San Diego, CA], and 2% glucose) at 30°C. *K. phaffii* transformants were selected in synthetic defined media lacking histidine (SD-his; 0.67% Difco YNB without amino acids, 0.069% CSM-his [Sunrise Science, San Diego, CA], and 2% glucose) at 30°C. *R. araucariae* and *K. marxianus* protoplasts were mixed with regeneration agar solution and incubated at 30°C for five to six days to allow for genome edits to occur. All transformations were performed in a minimum of three independent biological replicates.

#### Plasmid Construction

pSC012 (12), pIW601 (13), and pCRISPRpp (14) were used as the backbone plasmids to express Cas9 and sgRNAs in *Y. lipolytica, K. marxianus*, and *K. phaffii* genomes, respectively. Cloning sgRNAs in plasmid backbones was done following the method of Schwartz et al. (15). Each sgRNA was ordered as a primer from Integrated DNA Technology (IDT) with 20 bp homology up- and down-stream of the AvrII, PspXI, and BbvCI cut sites in pSC012, pIW601 and pCRISPRpp plasmids, respectively. 60-bp top and bottom strands were ordered and annealed together. The annealed strand and digested plasmid were assembled using Gibson Assembly in a 3:1 M ratio (insert:vector). The assembled product was then transformed directly into electrocompetent *E. coli* TOP10 cells. *E. coli* cultures were conducted in Luria-Bertani (LB) broth with either 100 mg/L ampicillin or 50 mg/L kanamycin at 37°C in 14 mL polypropylene tubes, at 225 rpm. Plasmid isolation was performed using the Zymo Research Plasmid Miniprep Kit. All plasmids, primers, and sgRNAs used in this work are listed in **Supplementary Table 2-4**.

#### Ribonucleoprotein Complex Assembly

For the generation of sgRNAs to form Cas9-RNP complexes, the EnGen® sgRNA Synthesis Kit, *S. pyogenes* (NEB #E3322), was used according to the manufacturer’s instructions. Target-specific DNA oligonucleotides were designed using the EnGen sgRNA Template Oligo Designer and checked for presence of a “G” at the 5’ end. If absent, a “G” was manually added. The resulting templates were then used for *in vitro* transcription with the EnGen® sgRNA Synthesis Kit. The transcribed sgRNA was treated with DNase I (NEB #E3322) and purified using the RNA Clean-Up Kit (NEB #T2040). Before RNP assembly, the sgRNA was denatured and refolded as described by Pohl et al. (16).

RNPs were assembled following the method of Kreuter et al. (17) with slight modifications. Cas9-RNPs were immediately assembled before experiments in a 50 µL reaction in buffer B (25 mM CaCl_2_, 10 mM Tris-HCl, 1 M sorbitol, pH 7.5) containing 5 µL 1X NEBuffer™ r3.1 (100 mM NaCl, 50 mM Tris-HCl, 10 mM MgCl_2_ and 100 µg/ml Recombinant Albumin, pH 7.9), and 1:1 molar ratio of sgRNA:Cas9 at 37°C water bath for 10 minutes.

#### CRISPR Genome Editing with ALLEGRO-Designed sgRNAs

We instructed ALLEGRO to design an sgRNA library with minimal size targeting a counter-selectable marker gene set (*Saccharomyces Cerevisiae* orthologs of *CAN1, FCY1, GAP1, LYS2, TRP1*, and *URA3*) across 2,263 fungal species using track *E*_*1*_, meaning that each gene in the set must be targeted at least once in each species (list of genes in **Supplementary Table 5**, and list of species in **Supplementary Data 1**). To validate the on-target activity of this sgRNA library, we conducted CRISPR-Cas9 gene knockout experiments across three different yeast species: *Kluyveromyces marxianus, Komagataella phaffii*, and *Yarrowia lipolytica*. Gene sets tested in each species included *CAN1, GAP1, LYS2* and *TRP1* in *Y. lipolytica, CAN1, FCY1*, and *URA3* in *K. marxianus*, and *CAN1* and *FCY1* in *K. phaffii*. The experimental workflow began with the selection of individual sgRNAs from the *E*_*1*_ library, targeting each set of auxotrophic genes in the desired genome and cloning into pIW601, pCRISPRpp, and pSC012 plasmid backbones, respectively, or assembled into Cas9-RNP complexes. For sgRNAs’ efficiency comparison, a set of active sgRNAs from previously designed and validated libraries (12, 14, 18) were selected as controls. All control sgRNAs are listed in **Supplementary Table 4**, and these were similarly cloned into respective plasmid backbones. These plasmids and RNP complexes were subsequently transformed into yeast cells and plated on synthetic defined media (SD) with and without the inhibitor of the targeted counter-selectable genes. To identify gene mutations, the chemical inhibitors specific to the auxotrophic genes were added to the SD plates (0.67% Difco YNB without amino acids, 0.079% CSM [Sunrise Science, San Diego, CA], 2% glucose and 2% agar). To control for false-positive colonies, yeast transformants or protoplasts were also transformed without the respective sgRNA, using either an empty vector or a Cas9-water complex. Colonies appearing on the chemical inhibitor plates were randomly picked and analyzed for frameshift or INDEL mutations. The list of chemical inhibitors associated to each gene is as follows: *CAN1*: L-canavanine, *FCY1*: 5-fluorocytosine, *GAP1*: L-Histidine, *LYS2*: a-aminoadipic acid, *TRP1*: 5-fluoroanthranilic, and *URA3*: 5-fluoroorotic acid. Selections for the loss of *CAN1, FCY1, GAP1, LYS2, TRP1* and *URA3* genes on SD plates containing-L-canavanine, 5-fluorocytosine, minimal L-proline+D-histidine medium (MPDHis), a-aminoadipic acid, 5-fluoroanthranilic acid and 5-fluoroorotic acid, respectively, was done following published methods (19–24).

#### Yeast Transformation and Screening

*Y. lipolytica* transformation was conducted by using a protocol described by Robertson et al. (12). Briefly, a single colony of the PO1f strain was grown in 2 mL of YPD liquid culture in a 14 mL culture tube at 30°C with shaking at 225 rpm for 18 hours (final OD600 ~30). A total of 350 µL of the culture was pelleted by centrifugation at 4,000 × g for 2 minutes and resuspended in 300 µL of transformation buffer (45% PEG 4000, 0.1 M lithium acetate (LiAc) and 100 mM dithiothreitol (DTT)). Next, 500 ng of plasmid DNA and 80 µg of 10 mg/mL Salmon sperm DNA (ssDNA, Agilent) were added to the mixture by thoroughly pipetting. Following incubation for 1 hour at 39°C, 1 mL of water was added, and the cells were pelleted and redistributed in 2 mL SD-ura. After 2 days of growth, the cells were plated at a 10-2 dilution on SD + L-canavanine (60 mg/L) and SD + MPDHis (10 mM) SD + a-aminoadipic acid (4.36 g/L), SD + 5-fluoroanthranilic (5-FAA) (1.5 g/L) to counter-select for *CAN1, GAP1, LYS2* and *TRP1* knockouts, respectively. Colonies were randomly picked from plates and the targeted sequence of the gene was PCR amplified. Gene knockout was verified by Sanger sequencing.

*K. marxianus* transformations were accomplished using a modified protocol as previously described (13). Briefly, a single colony of *K. marxianus* CBS6556 *ura3Δ* strain was inoculated in a 14 mL tube containing 2 mL YPD and grown overnight. This culture was used to inoculate a 250 mL baffled shake flask containing 50 mL of fresh YPD at a starting OD of 0.05, which was then grown for ~13 hours (final OD ~15-18). For each transformation, 7 × 10^8^ cells were collected and centrifuged at 4,000 × g for 1 minute at 4°C (1 OD600 = 1.4 × 10^7^ cells/mL). The cells were washed 3 times with 1 mL of the wash buffer (1 mM EDTA, 0.1 M LiAc) and then resuspended in 500 µL of the transformation buffer (38% PEG 4000, 1 M LiAc, 10 mM DTT, 10 mM tris-HCl, and 10 mM EDTA). Then, 4 µg of plasmid DNA and 150 µg ssDNA were added to the mixture, and the cells were incubated at room temperature for 15 minutes. Subsequently, the mixture was heat shocked in a 47°C water bath for 9 minutes. Following the heat shock, the cells were pelleted, the supernatant was removed, and cell pellets were resuspended into 2 mL SD-ura selective media for 2 days at 30°C. After 2 days, the cells were plated at a 10-2 dilution on SD + L-canavanine (50 mg/L) and SD + 5-fluorocytosine (1 mM) to counter-select for *CAN1* and *FCY1*, respectively. After 2-3 days of incubation at 30°C, colony PCR was performed to amplify the targeted region which was followed by Sanger sequencing to identify frameshift mutations leading to gene knockouts.

*K. phaffii* transformation was done using a previously described method (14). Briefly, 2 mL of YPD was inoculated with a single colony of *K. phaffii* GS115 *his4::CAS9* and grown in 2 mL YPD overnight. A total of 4 × 10^7^ cells were transferred to 150 mL of YPD in a 500 mL baffled shake flask and grown for ~14 hours when the final OD600 was ~ 1.8. 100 mL of cells were chilled on ice for 1.5 hours, washed three times with 1 M ice-cold sorbitol, incubated with 25 mL of pretreatment solution (0.1 M LiAc, 30 mM DTT, 0.6 M sorbitol, and 10 mM tris-HCl, pH 7.5) for 30 minutes at room temperature, and washed three more times with 1 M ice-cold sorbitol. For each transformation, 8 × 10^8^ cells were mixed with 2 µg of plasmid to a final volume of 80 µL, incubated on ice for 15 minutes, and pulsed at 1.5 kV with Bio-Rad MicroPulser Electroporator in an ice-cold 0.2-cm-gap cuvette (1 OD600 = 5 × 10^7^ cells/mL). Immediately after electroporation shock, 1 mL ice-cold solution of YPD and 1 M sorbitol was added to each cuvette. Cells were transferred to 1 mL YPD and 1 M sorbitol in 14 mL tubes, incubated for 3 hours at 30°C and 225 rpm for recovery, washed with 1 mL of room-temperature autoclaved water to dispose of the excess plasmid DNA in samples, and transferred to SD-his selective media for 3 days at 30°C. Cells reached to confluency after 3 days and were transferred into 2 mL fresh SD-his media with a starting OD600 of 1 to perform outgrowth experiments and were allowed to grow for 3 more days. After 6 days of growth, the cells were plated at OD600 = 1 on SD + L-canavanine (50 mg/L) and SD + 5-fluorocytosine (5-FC) (1 mM) to counter-select for *CAN1* and *FCY1*, respectively. Colonies were randomly picked from plates and the targeted sequence of the gene was PCR amplified, and gene knockout was verified by Sanger sequencing. All centrifugation was done at 3,000 × g for 5 minutes at 4°C.

#### Protoplasts Preparation and Transformation

Protoplasts of *K. marxianus* CBS6556 were prepared according to a previously described method (25) with slight modifications. Briefly, three fresh colonies of *K. marxianus* CBS6556 were scraped from the YPD plate and grew at 30°C overnight in a 250-mL flask containing 50 mL YPD medium. The following day, the overnight culture was transferred to a 50 mL fresh YPD media to prepare a day-2 culture with a starting OD600 of 0.05. The cells were grown at 30°C for 10-18 hours until OD600 of 6-8. From the culture, 25 mL was transferred into a 50-mL conical tube and then centrifuged at 1,000 × g for 5 minutes at 5°C. The supernatant was discarded, and the cells were resuspended in 30 mL of sterile water. Cells were centrifuged again at 3,000 × g for 5 minutes at 5°C, the supernatant was discarded, and cells were resuspended in 20 mL of citrate phosphate buffer (10 mM citrate phosphate, 1.5 M sorbitol, pH 6.8). Then, 70 µL of 10 mg m/L Zymolyase 20T solution (#E1005, Zymo Research), 25% w/v glycerol, 50 mM Tris-HCl, pH 7.5) and 40 *µ*L of β-mercaptoethanol (BME) were added. The samples were mixed gently vortexing and then incubated at 30°C for 45 minutes. Protoplast formation was monitored by measuring the OD600 of samples diluted 1:10 in 1.5 M sorbitol and comparing it to the OD600 in 5% SDS. Protoplast preparation was terminated for transformation when the ratio reached 5, as low transformation efficiency can result from either insufficient digestion or over-digestion. After incubation at 30°C, protoplasts were collected by centrifugation at 600 × g for 10 minutes at 5°C. After centrifugation, the supernatant was discarded, and the cells were resuspended in 10 mL of 1.5 M sorbitol with a wide-bore 5-mL pipette tip and then supplemented with more 1.5 M sorbitol to achieve a final volume of 30 mL. Next, protoplasts were centrifuged at 700 × g for 10 minutes at 5°C, and the washing step with 30 mL 1.5 M sorbitol was repeated once more. Finally, protoplasts were resuspended in STC solution (1.5 M sorbitol, 10 mM Tris-HCl, 50 mM CaCl_2_, pH 7.5) to a final concentration of 3-8 × 10^8^ cells/mL.

For the *K. marxianus* CBS6556 protoplast transformation, 200 µL of protoplast suspension was mixed with 50 µL of the RNP mix and Triton X-100 (0.006% [w/v] final concentration in transformation reaction) in a 14 mL tube to enhance cell membrane permeability (26). The mixture was incubated on ice for 25 minutes, as prolonged incubation time may improve editing efficiency (27). Then, protoplasts were supplemented with 1 mL of transformation buffer (40% PEG 4000, 10 mM Tris-HCl pH 7.5, and 50 mM CaCl_2_). Samples were mixed thoroughly by flipping over 5-10 times and then incubated on ice for 1 hour before centrifugation at 500 × g for 5 minutes at 5°C. Supernatants were discarded, and pellets were resuspended in 800 µL of STC solution gently and were mixed with protoplast regeneration agar, which consisted of 6.9 g/L yeast nitrogen base without amino acids, 2% glucose, 0.8 M sorbitol, 3% agar, 20 mg/L L-adenine hemisulfate, 20 mg/L L-arginine*·* HCl, 20 mg/L L-histidine*·* HCl, 30 mg/L L-Isoleucine, 100 mg/L L-leucine, 30 mg/L L-lysine*·* HCl, 20 mg/L L-methionine, 50 mg/L L-phenylalanine, 200 mg/L L-threonine, 20 mg/L L-tryptophan, 30 mg/L L-tyrosine, 20 mg/L uracil, and 150 mg/L L-valine. The regeneration agar was melted and equilibrated at 47°C in advance. Transformed protoplasts were mixed with 10 mL of regeneration agar and 5-fluoroorotic acid (5-FOA, 1 g/L) in a 15-mL falcon tube. The tubes were flipped three to five times to mix the contents, and the mixture was poured onto the surface of SD + 5-FOA plates (1 g/L). Plates were incubated at 30°C until transformants formed, and then colony PCR and Sanger sequencing was performed to find frameshift mutation gene knockouts.

Protoplast transformation was also performed for the *R. araucariae* NRRL Y-17376 strain. We used the frozen protoplast protocol following a previously described method (28) with slight modifications. Briefly, single colonies of NRRL Y-17376 were separately inoculated in 50 mL of YPD medium at 30°C for 15 hours. Cells were harvested at 3,000 rpm for 10 minutes and suspended in 20 mL autoclaved H_2_O. Cells were harvested again, gently resuspended in 10 mL 1 M sorbitol, and subsequently harvested. Finally, the cells were suspended in 10 mL SCEM (1 M sorbitol, 0.1 M sodium citrate, 10 mM EDTA, 30 mM 2-mercaptoethanol, pH 5.8). Then, cells were mixed with 40 µl of lyticase solution (25,000 U/mL, Sigma Aldrich) and incubated at 30°C for 1 hour. Following lyticase digestion, the cells were harvested by centrifugation at 1200 × g for 10 minutes and resuspended in SCEM to a final concentration of 10^9^ cells/mL. Subsequently, 0.5 mL of lytic enzyme solution (1.5% [w/v] Zymolyase Ultra 2000U) was added to 1 mL of the cell suspension, and the mixture was incubated overnight at 30°C. Next, the cells were centrifuged gently at 300 × g for 5 minutes at 4°C in round-bottom plastic tubes and suspended in 10 mL 1 M sorbitol by gently tapping the tube. Then cells were centrifuged at 300 × g for 5 minutes. This procedure was repeated two more times to remove lyticase thoroughly, with the supernatant being discarded at each step. Finally, cells were suspended in 2 mL CaST solution (1 M sorbitol, 10 mM Tris-HCl, 10 mM CaCl_2_, pH 7.5) along with 2 mL of cell storage solution, and the protoplasts were stored at −80°C.

For the *R. araucariae* protoplast transformation, 200 µL of protoplast suspension was mixed with 150 µL of the RNP mix and Triton X-100 (0.006% [w/v] final concentration in transformation reaction) in a 2 mL tube. The mixture was kept on ice for 25 minutes. Then, the cells were transferred to an ice-cold 0.2-cm-gap aluminum cuvette and electroporation was performed (1.2 kV or 400 V, 400 Ω and 25 µF capacitance), using a Bio-Rad MicroPulser Electroporator. After electroporation, the cells were resuspended in 1 mL ice-cold YPD and transferred into new tubes on ice for 30 minutes, and then incubated at 30°C for 4 hours. Protoplasts were mixed with protoplast regeneration agar consisting contained 5 g/L glucose, 4 g/L KH_2_PO_4_, 0.5 g/L Na_2_HPO_4_, 3.0 g/L NH_4_C_1_, 0.5 g/L NaCl, 0.4 g/L MgSO_4_*·*7H_2_O, 0.01 g/L CaCl_2_*·* 2H_2_O, 0.008 g/L FeCl_3_*·* 6H_2_O, 0.0001 g/L ZnSO_4_*·* 7H_2_O, 0.069% CSM, pH 5.5, supplemented with 1.5 M sorbitol and 2.5% agar. The regeneration agar was melted and equilibrated at 47°C in advance. Transformed protoplasts were mixed with 10 mL of regeneration agar, respective concentrations of a-aminoadipic acid and 5-fluorocytosine in a 15-mL Falcon tube. The tubes were flipped over three to five times quickly and then the mixture was poured on the top of SD + a-aminoadipic acid, and SD + 5-fluorocytosine plates. Plates were incubated at 30°C until transformants formed, and then colony PCR, and Sanger sequencing was performed to find frameshift mutation gene knockouts.

## Results

To create minimal Cas9 sgRNA libraries using ALLEGRO’s two design tracks across the fungal kingdom, we downloaded the genomes, protein sequences, and GFF files for 2,263 fungal species from NCBI (29), FungiDB (30), EnsemblFungi (31, 32), and MycoCosm (33), the list of which is provided in **Supplementary Data 1**. We extracted the transcriptome of each genome using the corresponding GFF and recorded the intron-exon boundaries. An illustrative tree that shows the composition of the data set is provided in **Supplementary Figure 2**. The scripts and datasets, along with the documentation on how we carried out this step can be found at https://github.com/ucrbioinfo/fugue.

**Fig. 2.**
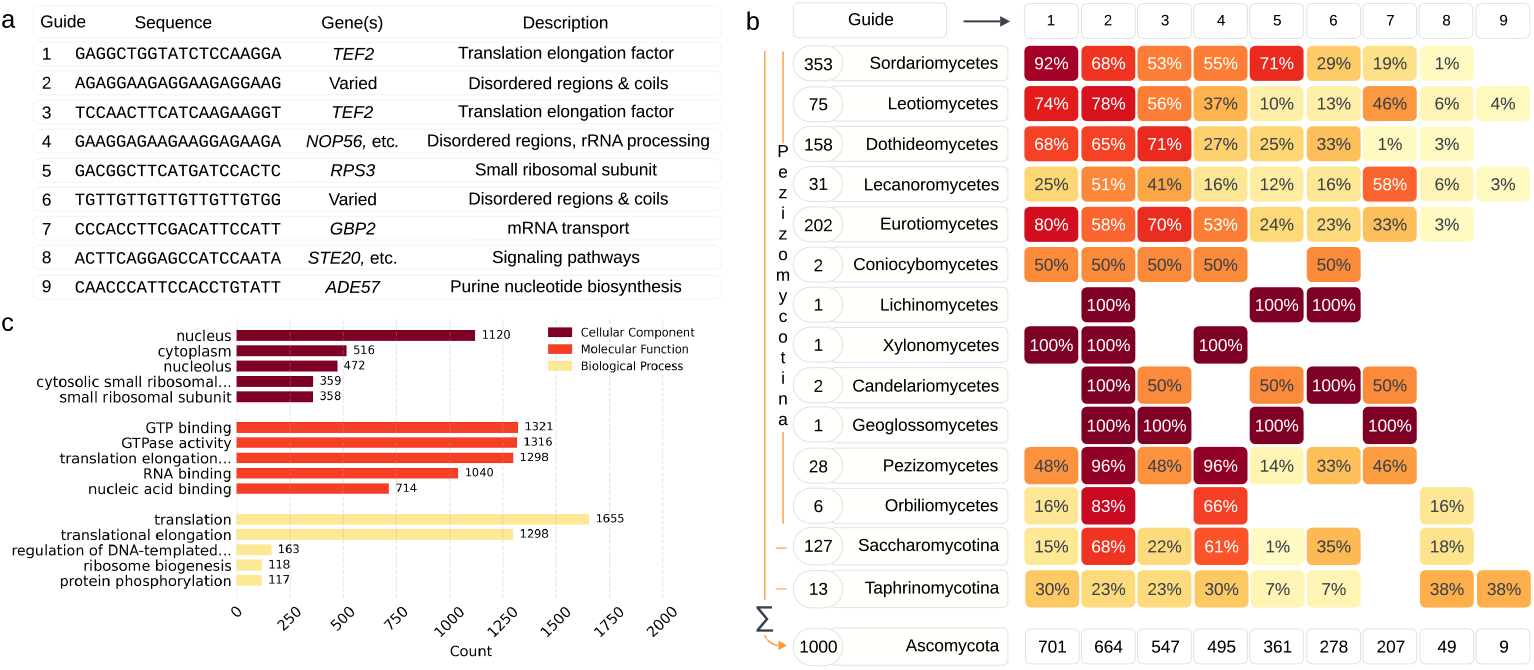
ALLEGRO scales to thousands of species for genome-wide library design. **(a)** ALLEGRO produced a library of nine guides that target at least one gene in 1,000 *Ascomycota* species. The sequence of the guides in the library with a brief description of their functions are shown. **(b)** An illustration of the 1,000 input *Ascomycota*, in which the numbers to the left of each species’ name indicate how many species from that group are included in the dataset. Percentages in each column indicate the portion of each species group targeted by each guide. The numbers in the bottom row indicate the total number of targets by each guide. **(c)** The top five Gene Ontology (GO) terms and their counts for each functional category for genes targeted by the nine guides, with each category representing biological process, molecular function, or cellular component.

### Nine Guides Target the Entire Transcriptome of One Thousand *Ascomycota*

In this first experiment, we wanted to stress-test ALLEGRO’s scalability to handle the entire transcriptome (i.e., all genes) for up to one thousand fungal species. To this end, we used ALLEGRO to calculate the smallest library of sgRNAs that would make at least one cut anywhere in the coding genes of all 1,000 *Ascomycota* (the full list of species is provided in **Supplementary Data 2**). Configured with track *A*_*1*_, and given the entire transcriptome of 1,000 *Ascomycota* species, ALLEGRO considers more than a billion guides. This high number of guides remain in the set even after removing guides containing homopolymers of length four or higher and guides with a GC content outside of the 40%-60% range, factors which are known to affect sgRNA efficiency (34, 35). ALLEGRO, however, can safely ignore millions of potential guides without affecting the quality or size of the final library by using a heuristic which we call the *redundancy threshold r*. For example, in *A*_*m*_ we can safely discard all but *m* guides that target up to the exact same *r* species without affecting the final size of the library. This concept extends to track *E*_*m*_ as well. In the *E*_*m*_ case, we discard all but *m* guides that target the exact same set of genes of size up to *r* (if configured to use scores, we would keep the best *m* guides). We outline a more detailed description of this heuristic in **Supplementary Notes Section 1.6**. Using a redundancy threshold set to 1,000 (equal to the number of input species for *A*_*1*_), the candidate set of guides was reduced from more than a billion to about 8.2 million. ALLEGRO solved the linear program with 8.2 million variables in about 13.5 hours using about 400 GB of RAM.

ALLEGRO produced a surprisingly small solution for the *A*_*1*_ track for the 1,000 *Ascomycota*. A library of only nine guides can target at least one gene in every species. Each guide, its sequence, most targeted gene, and description of gene function is shown in Figure 2a. Figure 2b illustrates the 1,000 *Ascomycota*, and the number of species in each group targeted by each guide. The first few guides target the majority of the species, while the last two guides are needed to achieve the full coverage of the 1,000 species, as counted in the bottom row of Figure 2b.

Given the small number of guides needed to cover all species, we examined whether these guides were more likely to target species that are closely related in terms of evolution. Guide 1, which targets exclusively *TEF2* orthologs, affects mostly fungal species in the *Pezizomycotina* family, except for those in *Lecanoromycetes* and *Orbiliomycetes*. In fact, 92% of all *Sordariomycetes* and 80% of *Eurotiomycetes* in the dataset are targeted by this guide, i.e., they share this exact DNA sequence in their *TEF2* orthologs. Guides 2 and 4 target almost all *Pezizomycetes* and more than half of *Sordariomycetes, Eurotiomycetes*, and *Saccharomycotina*. Guide 7 mostly targets the *GBP2* gene in almost half of the species from *Leotiomycetes, Lecanoromycetes*, and *Pezizomycetes*, and about one-third of those in *Eurotiomycetes*, with fewer targets in *Sordariomycetes*. The majority of the targets of Guide 5, which targets the orthologs of *RPS3*, are in *Sordariomycetes*, with a quarter of *Dothideomycetes* and *Eurotiomycetes* also targeted. Guides 8 and 9, having the fewest targets, cut mostly in *Saccharomycotina* and *Taphrinomycotina*, the two subdivisions in this dataset separate from the rest of the *Pezizomycotina*. **Supplementary Figure 3** shows the number of species targeted for any subset of these guides. The upset plot shows a strong overlap between the sets: for instance, while Guide 1 targets 701 species overall, it targets only 17 species that no other guide targets.

**Fig. 3.**
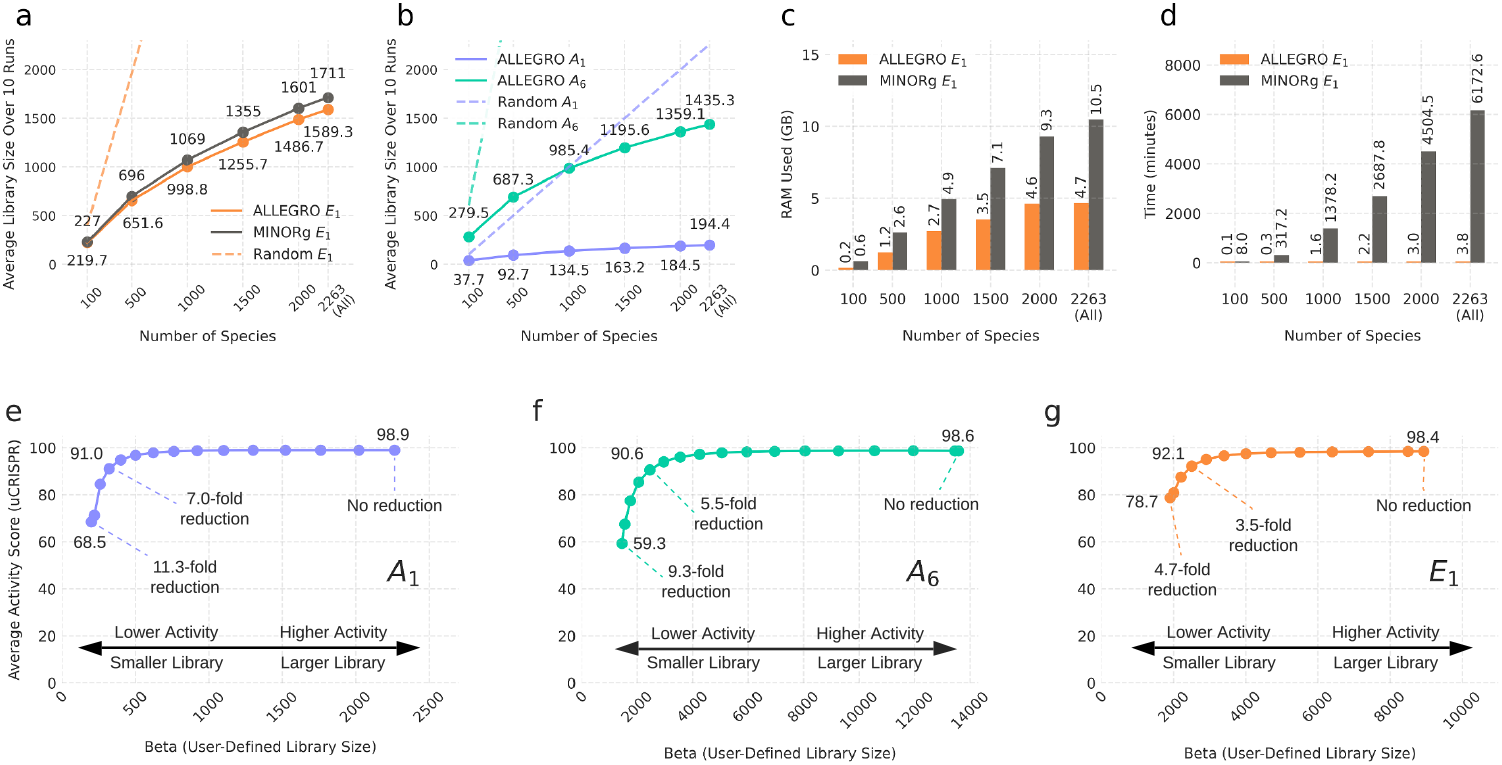
ALLEGRO enables the design of fungal kingdom-wide sgRNAs. **(a)** Library sizes generated by MINORg and ALLEGRO track *E*_*1*_ as a function of the number of species. In this comparison, we allowed guides crossing intron-exon boundaries in ALLEGRO as it is done in MINORg, and configured ALLEGRO with a redundancy threshold set to the maximum number of genes available. Random *E*_*1*_ refers to a naive sgRNA selection strategy where a random guide per gene is chosen. **(b)** Library sizes for ALLEGRO tracks *A*_*1*_ and *A*_*6*_ as a function of input size. Here, the redundancy threshold was set to the maximum number of input species. **(c)** The amount of memory used (in gigabytes) by MINORg and ALLEGRO (track *E*_*1*_) as a function of the number of input species **(d)** Running time (in minutes) of ALLEGRO (track *E*_*1*_) and MINORg as a function of the number of input species. **(e-g)** The beta parameter fine-tunes the trade-off between guide library size and the guides’ average scores.

Figure 2c illustrates a Gene Ontology (GO) enrichment analysis for genes targeted by the library guides. Notably, Guides 1 and 3 exclusively target orthologs of *TEF2*, which, according to (36), is a conserved gene crucial for translation elongation and accuracy. These guides specifically target the “Translation factor GTPase” family, focusing on the “Translational (tr)-type GTP-binding domain.” Guide 4 primarily targets disordered regions of orthologs such as *NOP56, NOP6, NHP2, CBF5*, and *DBP3*, which are involved in ribosome biogenesis and rRNA processing, localized to the nucleus (37, 38). Guide 5 exclusively targets orthologs of *RPS3*, an essential gene encoding proteins of the cytosolic small ribosomal subunit (40S), which localize to both the nucleus and cytosol (39). Guide 7 targets orthologs of *GBP2* and its paralog *HRB1*. These genes encode RNA-binding proteins involved in mRNA transport from the nucleus to the cytoplasm, playing a key role in mRNA export, especially under stress conditions. They are crucial surveillance factors for the selective export of spliced mRNAs in yeast, interacting with the nuclear RNA export factor *MEX67*. While non-essential, *GBP2* and *HRB1* are important for efficient RNA processing and stress response (40). Guide 8 targets orthologs such as *STE20, KIC1*, and *CLA4*, members of the PAK (p21-activated kinase) family involved in signaling pathways regulating cell growth, polarity, and responses to environmental cues. *STE20* plays a key role in the mating and filamentous growth pathways, activated by GTP-bound *CDC42*, a small GTPase, to transduce signals for mating and growth (41). *KIC1* is involved in cell wall integrity and polarity via the RAM pathway, regulating cell separation and polarized growth (42). *CLA4*, also *CDC42*-activated, is essential for cytokinesis and polarity maintenance (43). Lastly, guide 9 targets orthologs of *ADE57*, a gene involved in the *de novo* purine nucleotide biosynthesis pathway. This bifunctional enzyme, comprising aminoimidazole ribotide synthase and glycinamide ribotide synthase activities, is essential for purine synthesis (44).

Further analysis of the target sites revealed that Guides 2 and 6 do not target a specific gene or gene family but instead a diverse set of genes, predicted to target disordered protein regions (**Supplementary Data 3**). These disordered regions contain shared sequences across many genes, explaining the broad targeting of these guides. Additionally, guides 2, 4, and 6 exhibit significant similarity and high levels of repetition. According to (45), simple amino acid sequences are the most frequently occurring protein fragments in Saccharomyces cerevisiae. Moreover, some studies (see e.g., (46–48)) suggest that intrinsically disordered protein regions are usually composed of compositionally-biased and low-complexity sequences, corroborating our findings.

### ALLEGRO Scales to Auxotrophic Genes in Over Two Thousand Fungal Species

One use of ALLERGO is designing CRISPR guides that target many different species across a large number of organisms from the same kingdom. In this section, we demonstrate ALLEGRO’s capability in designing sgRNA libraries targeting auxotrophic genes in various combinations (tracks) for 2,263 fungal species where track *A*_*1*_ designs a library of sgRNAs targeting any of the genes of interest at least once. Track *A*_*6*_ is similar to *A*_*1*_ but targets anywhere across the six genes at least six times. Track *E*_*1*_ targets each individual gene at least once. By leveraging the counter-selectability of these genes and the sgRNA libraries designed by ALLEGRO, we aim to facilitate the domestication and onboarding of new fungal species.

We grouped the protein-coding genes for 2,263 fungal species representing isolates from across the fungal kingdom into orthogroups using DIAMOND (10). We used the protein sequences of *CAN1, FCY1, GAP1, LYS2, TRP1*, and *URA3* in *S. cerevisiae* to identify the orthogroups for these genes. All input species have at least one gene in the orthogroup with at least 30% sequence identity to those in *S. cerevisiae*. We chose this level of identity based on prior studies on structural similarity (48–50).

### Library Size as a Function of the Number of Species

The objective of this experiment was to study the size of ALLEGRO’s libraries for tracks *A*_*1*_, *A*_*6*_, and *E*_*1*_ as a function of the number of input species. We also wanted to compare the size of the library produced by ALLEGRO in track *E*_*1*_ to the size of the libraries produced by MINORg. Both ALLEGRO and MINORg were executed on the same inputs and used the same parameters, including the use of multi-threading, not checking for off-targets, allowing no mismatches, and allowing only guides with a GC content between 30-70%. The inputs and commands used to run MINORg, and the configuration files for ALLEGRO can be found in **Supplementary Data 4**.

We randomly sampled *x* species from the pool of 2,263, each with at least one gene out of the six auxotrophy genes. For each choice of *x* = *{*100, 500, 1000, 1500, 2263*}*, we created ten samples of *x* species to reduce the sampling bias. ALLEGRO was executed on ten samples for all tracks, while MINORg was run on a single sample due to its high running time. To ensure fairness, for each species set, the sample where ALLEGRO’s library size was closest to the average across the ten samples was selected as the input for MINORg. This approach was used to minimize variability and provide a consistent benchmark for comparing the performance of both tools. For both tools, we recorded the size of the library as well as the time/memory used. The results presented in Figure 3a illustrate the average library sizes across the ten runs for ALLEGRO, and the library sizes generated by MINORg. **Supplementary Figure 4** shows the means and standard deviations of ALLEGRO’s library sizes.

**Fig. 4.**
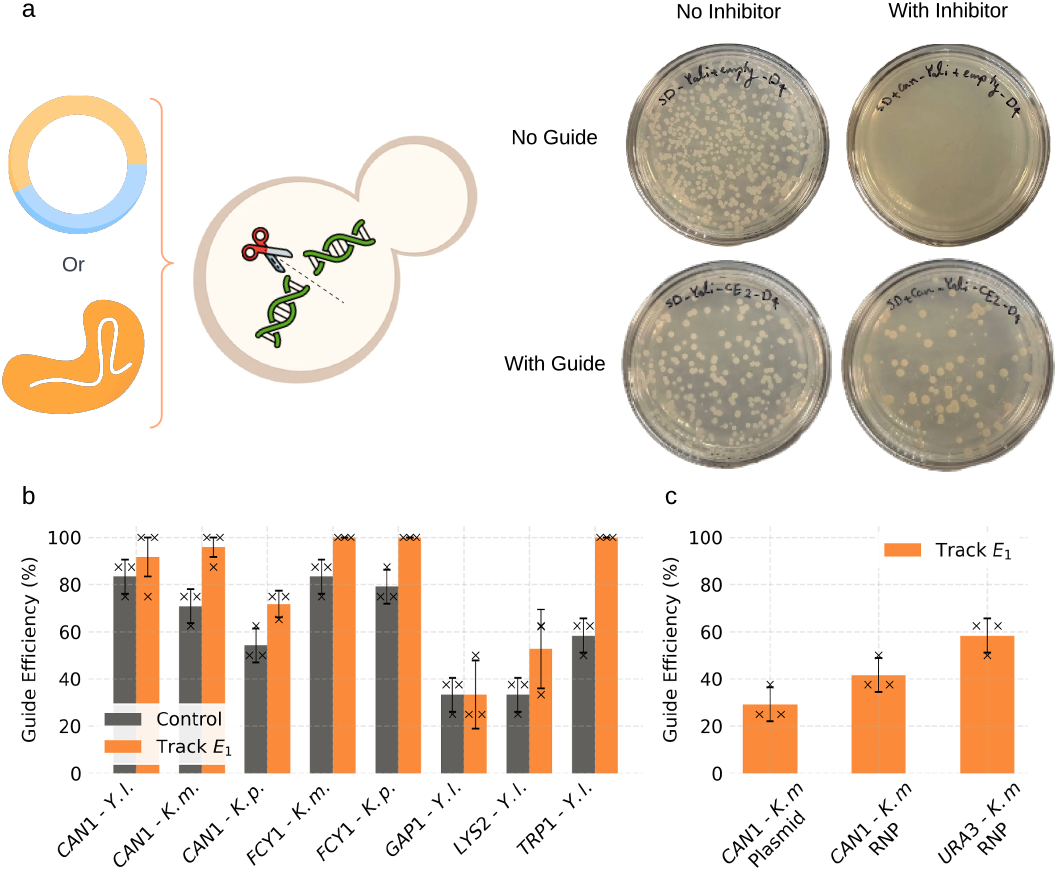
Experimental validation for ALLEGRO’s track *E*_*1*_. **(a)** CRISPR-Cas9 knockout screening. Track *E*_*1*_ and control sgRNAs targeting auxotrophic genes are cloned into plasmid backbones and transformed into the yeast of interest. Transformants are plated on standard control media (SD) and inhibitor-containing media, with experiments conducted in triplicate. Colonies growing on inhibitor media are validated using colony PCR. Selected images are an illustration of control and sample plates for targeting *CAN1* gene in *Yarrowia lipolytica* **(b)** CRISPR-Cas9 cutting efficiency scores of ALLEGRO-designed sgRNAs targeting auxotrophic genes in *Yarrowia lipolytica* (Y. l), *Kluyveromyces marxianus* (K. m.) and *Komagataella phaffii* (K. p.). From each of the three biological replicates, eight colonies appearing on inhibitor-containing plates are randomly selected and sequence analyzed to confirm the presence of mutations. **(c)** CRISPR-Cas9 cutting efficiency scores of ALLEGRO-designed sgRNAs targeting *CAN1* and *URA3* genes in *K. marxianus* genome using protoplast transformation with plasmid or Cas9-RNP complexes. The error bars represent the standard deviation and bars represent the mean. Individual data points for all experiments are also shown.

Figure 3a shows the size of the library as a function of the number of species *x*. To establish a baseline, a naive algorithm, Random *E*_*1*_, was also included. Random *E*_*1*_ generates libraries by selecting one random sgRNA for each targeted gene in each species, ultimately dropping any potential duplicates. ALLEGRO’s library size is smaller than MINORg’s for all choices of *x*. Figure 3b shows the average sizes of the *A*_*1*_ and *E*_*6*_ libraries as a function of the number of species *x* over 10 runs. Figure 3c shows the peak RAM usage of MINORg and ALLEGRO in gigabytes. ALLEGRO uses almost half as much memory as MINORg given the largest available dataset. Additionally, ALLEGRO completes its tasks in minutes while MINORg may take up to multiple days to complete the calculation (Figure 3d).

In the next ensemble of experiments, we added two extra criteria to our ALLEGRO libraries where we (i) allowed up to a single mismatch between the sgRNA sequence and its target in the PAM-distal region, and (ii) did not allow sgRNAs that crossed an intron-exon boundary. We allowed criteria (i) because we found that the library size can be significantly reduced if a mismatch is allowed. According to previous studies (1, 51–54), a single mutation in the 8 bases of the 5’ seed-distal region is tolerated, while mismatches in the first 12 bases upstream of the 3’-NGG abolish Cas9 cleavage. Further, criteria (ii) must be considered because an sgRNA target identified within a protein-coding sequence and spanning through a splice site may be ineffective in cutting its intended target in the genome due to sequence separation. Designing a library to target each auxotrophy gene across all species at least once (track *E*_*1*_) would trivially require 13,578 sgRNAs (2,263 species × 6 genes). However, with ALLEGRO’s *E*_*1*_, this number can be drastically reduced to 1,809 sgRNAs—a reduction of 86%—if mismatches are not allowed, and to 1,485 sgRNAs—a reduction of 89%—if mismatches are allowed. The library sizes for tracks *A*_*1*_, *A*_*6*_ and *E*_*1*_ for ALLEGRO can be found in Table 1.

### Trade-off Between Set Size and Guide Activity

In the experiments carried out so far we have not considered guide activity. The objective of minimizing the size of the library competes against the objective of maximizing the predicted activity scores for the library. Allowing the library size to grow gives ALLEGRO the flexibility to choose low activity guides that cover more targets and vice versa. This trade-off is controlled by the *beta* parameter, the library size budget. By setting beta to be higher than the smallest library size, we allow for the average predicted activity scores for the library to increase.

This trade-off is illustrated in our experiments for tracks *A*_*1*_, *A*_*6*_, and *E*_*1*_ on the six auxotrophic marker genes *CAN1, FCY1, GAP1, LYS2, TRP1*, and *URA3* from 2,263 fungal species (Figures 3e, f, and g). For track *A*_*1*_ the largest possible beta is 2,263 when choosing the highest activity guide per species. As beta decreases, the library size becomes smaller at the expense of the average activity score. With a beta of 320, the library contains 320 guides (a 7.0 fold reduction) and the average activity score for the library is still quite high at 91.0. The smallest library possible is for beta = 212 which gives a 11.3-fold reduction from the original library size, but the average predicted score is low at 68.7. We observe a similar trade-off for track *A*_*6*_ (Figure 3f) where for beta = 2,450, the library 5.5-fold reduced from the maximum size while maintaining a high average guide activity score of 90.6. The smallest beta for this experiment is 1,450 (a 9.3-fold reduction) with a low average guide score of 59.2. A similar trend is observed in track *E*_*1*_ (Figure 3g) where with a 3.5-fold reduction at beta = 2,500, the average guide score is 92.1. With a 4.7-fold reduction at beta = 1,900, we have a library with an average guide score of 78.7. For *A*_*6*_ the largest beta is 2,263 × 6 = 12,330 where each species may be targeted six times. For *E*_*1*_ the largest beta is 8,932 which is the total number of available orthologous genes across the 2,263 species. All data used to produce **Figure 3** may be found in **Supplementary Data 4**.

### Supplementary Data 4

#### Track E_1_ Achieves Validation-Grade Efficiency in Fungi

To demonstrate ALLEGRO’s capability in designing active sgRNAs, we experimentally validated the targeting efficiency of sgRNAs designed for the track *E*_*1*_ library that target the counter-selectable marker gene set (*CAN1, FCY1, GAP1, LYS2, TRP1*, and *URA3*). In these experiments, three yeast species—*Kluyveromyces marxianus, Komagataella phaffii*, and *Yarrowia lipolytica*—were transformed with plasmid DNA consisting of Cas9 and sgRNA expression cassettes optimized for each species. Colonies on selective media with chemical inhibitors, which required gene disruption for growth, were analyzed for knockouts (Figure 4a). As shown in Figure 4b, ALLEGRO-designed sgRNAs exhibited high targeting efficiency, achieving 90-100% cutting efficiency when targeting *CAN1* and *TRP1* in *Y. lipolytica* and *CAN1* and *FCY1* in *K. marxianus*. Notably, ALLEGRO designed guides outperformed control sgRNAs across these targets. Additional data regarding the number of colonies observed on both control and selective media is shown in **Supplementary Figure 5-7**. Aside from targeting specific genes in *K. marxianus, K. phaffii*, and *Y. lipolytica*, **Supplementary Data 5** provides a phylogenetic analysis of the broader fungal groups targeted by track *E*_*1*_ sgRNAs validated in these species. This analysis highlights the evolutionary relationships and demonstrates the broader applicability of the designed sgRNAs across related fungal groups.

**Fig. 5.**
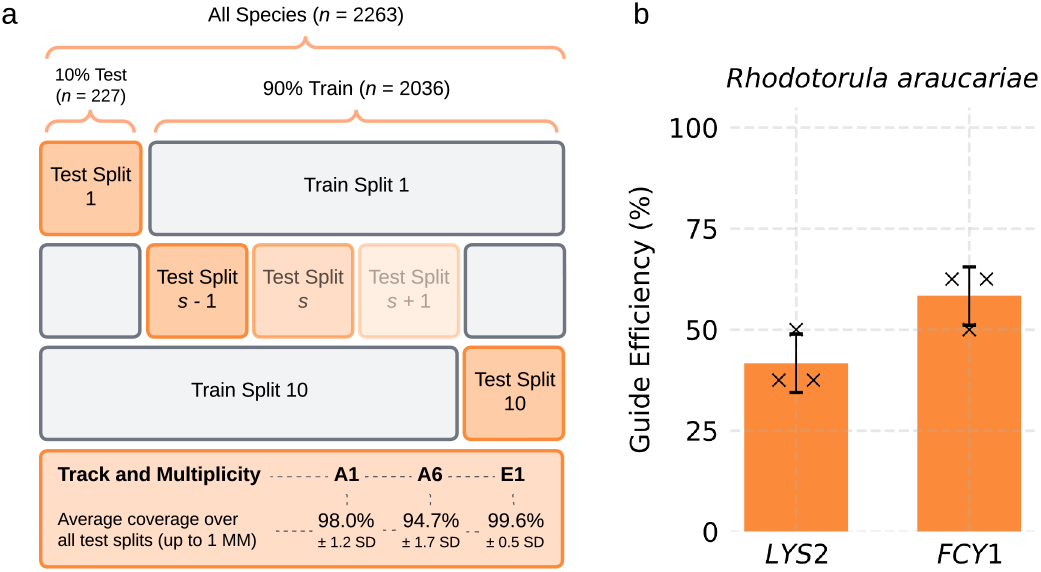
Robustness Analysis of ALLEGRO. **(a)** Cross-species validation results. 10% of the species were excluded as a test set, and the pipeline was trained on the remaining 90%. The sgRNA library size and average coverage were calculated for the excluded test splits. The results demonstrate high coverage and highlight ALLEGRO’s ability to design efficient sgRNAs across diverse species. **(b)** ALLEGRO’s performance on fungal species not included in the training set. sgRNAs from track *E*_*1*_ library were tested for targeting *FCY1* and *LYS2* in *R. araucariae*.

The next validation experiments assessed the broader applicability of ALLEGRO designed sgRNAs to enable genome editing in fungi using a universal transformation method. This approach combined Cas9-RNP complexes with protoplast transformation. Initially, we demonstrated the feasibility of protoplast transformation in the *K. marxianus* genome by targeting the *CAN1* gene with plasmid DNA. We then showed that this method, coupled with RNP complexes, efficiently targets the *CAN1* gene (Figure 4c). Furthermore, we demonstrated that Cas9 RNPs complexed with ALLEGRO-designed sgRNAs efficiently knockout the *URA3* gene with a cutting efficiency of ~60% in *K. marxianus*. **Supplementary Figure 8** provides additional data on the number of colonies observed on both control and selective media for *K. marxianus* protoplast transformation targeting the *CAN1* gene.

### Kingdom-Wide Guide Design with ALLEGRO

One of the goals of our work is to design a set of universal guides that would target a given set of genes of interest in every fungal species. Since our genomic knowledge of fungi is limited to the *≈*2,000 species whose genome has been sequenced, in this experiment we attempt to quantify how guides designed for a subset of species would also work for other species not included in ALLEGRO’s training set.

To answer this question, we conducted 10-fold cross-validation experiments for tracks *A*_*1*_, *A*_*6*_, and *E*_*1*_ on the six counter-selectable marker genes *CAN1, FCY1, GAP1, LYS2, TRP1*, and *URA3*. In each calculation, we split the 2,263 species into 90% training (2,036 species) and 10% testing (227 species), as illustrated in Figure 5a. We ran ALLEGRO on the training set of each split and determined whether the guides in the library would also cut these six genes in the 10% test species. Guide design was successful if the guide matched the gene sequence with up to one mismatch in the seed-distal region. We measured the performance of ALLEGRO on each test set by counting the number of species that (i) would have every gene cut somewhere at least once (track *E*_*1*_), (ii) would have any of the six genes cut somewhere at least once (track *A*_*1*_), or (iii) would have the set of genes cut somewhere at least six times (track *A*_*6*_). If a sufficient number of cuts was produced in a species to satisfy the track conditions, we said that the species was covered.

We averaged the number of test species covered by ALLEGRO’s libraries over the 10-fold cross-validation experiment. We found that track *E*_*1*_ produces libraries that on average cover 99.6% of the test (unseen) species. For track *A*_*1*_, ALLEGRO’s libraries cover on average 98.0% of the test species. The least generalizable library is track *A*_*1*_. The libraries produced by ALLEGRO for *A*_*6*_ cover 94.7% of all test species. The bottom part of Figure 5a summarizes these findings. This experiment suggests that there is a high likelihood of obtaining a universal set of guides for a specific set of genes of interest, in particular when the set of species sequenced will be more representative of the diversity of fungi on the planet. Scripts to reproduce the cross-validation experiment are included in **Supplementary Data 6**.

To test ALLEGRO’s capability in designing Cas9-sgRNAs for a fungi species not included in the training set, we set out to create auxotrophic strains of *R. araucariae*. A BLAST+ (ncbi-blast-2.16.0, (55)) analysis was performed using the *R. araucariae* genome and the track *E*_*1*_ sgRNA library. The analysis revealed that the library successfully targeted *LYS2, URA3*, and *FCY1* in *R. araucariae*. No orthologs for *CAN1* and *TRP1* were identified. Additionally, a BLAST search confirmed that no sgRNAs in the *E*_*1*_ library were available to target *GAP1*.

Based on these findings, we proceeded to validate sgRNAs from the *E*_*1*_ library that specifically target the *FCY1* and *LYS2* genes (Figure 5b). The sgRNAs were validated through a Cas9-knockout screening using protoplast transformation and RNPs. The list of validated sgRNAs is provided in the **Supplementary Table 4**. Further data regarding the number of transformants under control and selective conditions for *R. araucariae* transformation is presented in **Supplementary Figure 9**. These results confirm ALLEGRO’s capability to design active and efficient sgRNAs for genome editing in novel fungal species.

## Discussion

We introduced a CRISPR guide design tool called ALLEGRO capable of efficiently designing guide libraries with minimal cardinality and scaling to two thousand species. ALLEGRO leverages advanced combinatorial optimization tools to find the smallest set of guides that target a set of genes of interest. Previous research has concentrated on designing guide libraries for a small set of species and genes. Based on our experiments, these tools struggle with large datasets, either fail to produce any results, or consume substantial time and computational resources and often generate outputs that are sub-optimal.

As a proof of scalability, we showed that ALLEGRO can design guides for all the genes (i.e., the full transcriptome) for a thousand *Ascomycota* species. Due to the infeasibility of storing more than a billion candidate guides in main memory, we developed new methods to disregard guides without affecting the optimality of the solution. Using the new approach, ALLEGRO identified just nine guides capable of targeting anywhere across the transcriptome of the 1,000 input species.

We also carried out extensive experiments on a set of six auxotrophic genes on more than two thousand fungal species. We showed that ALLEGRO produces a smaller guide library than MINORg, requires less RAM, and is several orders of magnitude faster. We also showed that the size of the required guide library for tracks *A*_*1*_, *A*_*6*_, and *E*_*1*_ tends to plateau as the number of input species increases, which is a required feature of a universal library that would target all the species within a kingdom. To this end, we performed several cross-validation experiments, designing guides for a portion of the input and testing them on a small subset of unseen species. By allowing a single base mismatch in the seed-distal region of the guide and its potential targets, we showed that we could meet the requirements for each track in over ~95% of the held-out species on average.

A unique feature of ALLEGRO is that it can incorporate the predicted guide activity scores to design the optimal library. We showed that while the smallest guide library is likely to contain low-activity guides, a set with higher predicted activity may be achieved by allowing ALLEGRO to select a slightly larger number of guides. While ALLEGRO is designed to be fast and flexible, large experiments such as our full transcriptome experiment over a thousand species, still require a significant amount of primary memory. To address this issue, new heuristics must be developed to discard guides without compromising the size of the final library. In addition, new linear programming solvers might yield better results in terms of library size, processing time, and memory usage.

We experimentally validated ALLEGRO’s ability to design highly efficient and precise sgRNAs for genetic engineering through CRISPR-Cas9 knockout screens. Our results confirmed successful knockouts of auxotrophic genes in multiple industrially-relevant yeast species, including *Kluyveromyces marxianus, Komagataella phaffii, and Yarrowia lipolytica*, demonstrating the high efficiency and on-target activity of ALLEGRO-generated sgRNAs. Furthermore, computational cross-validation, and successful experimental validation in *Rhodotorula araucariae*, a species not included in the initial input set, underscores ALLEGRO’s broad applicability for genome editing across diverse and novel fungal species. These findings establish ALLEGRO as a powerful, adaptable tool for precision genome editing across entire biological kingdoms.

## Data Availability

ALLEGRO is freely available on GitHub at https://github.com/ucrbioinfo/allegro and on Zenodo at 10.5281/zenodo.14768175. ALLEGRO’s documentation is provided on its GitHub Wiki page. All input data used in this work can be obtained from ALLEGRO’s repository and the **Supplementary Data**. For reference, we provide the pre-processing procedures for acquiring the input data in FUGUE’s repository https://github.com/ucrbioinfo/fugue. *R. araucariae* genome is deposited on NCBI BioProject through accession PRJNA895933.

## Supplementary Data

### Supplementary Data 1

ALLEGRO input for 2,263 fungal species is on Zenodo at fourdbs_input_species.csv. The actual genomic files to which this manifest points can be found on Zenodo at 2263_ortho_input.zip.

### Supplementary Data 2

ALLEGRO input manifest for 1,000 *Ascomycota* species is on Zenodo at ascomycota_input_species.csv.

### Supplementary Data 3

PDF for 1,000 *Ascomycota* nine guides, their GO terms, and target protein domains available on bioR*χ*iv.

### Supplementary Data 4

ALLEGRO and MINORg (version v0.2.3.4alpha0) commands, inputs, and outputs used to create Figure 3 are on Zenodo at allegro_fig3_data.zip.

### Supplementary Data 5

PDF for phylogenetic tree of the broader fungal groups targeted by track *E*_*1*_ sgRNAs validated in *K. marxianus, K. phaffii*, and *Y. lipolytica* available on bioR*χ*iv.

### Supplementary Data 6

ALLEGRO cross-validation scripts, inputs, and outputs are on Zenodo at allegro_fig5_data.zip.

## Acknowledgements

IW and SL conceived and supervised the project. AM developed ALLE**G**R**O** and conducted the computational experiments. RGN performed the experimental validations. AT assisted in experimental methodology, writing, reviewing, and editing. **J**ES and XL prov**i**ded the ***R***. *araucariae* genome sequence, and the strain was provided by the USDA-ARS Culture Collection (NRRL). **A**ll authors wrote and edited the manuscript.

