## Supplementary Materials for "Kingdom-Wide CRISPR Guide Design with ALLEGRO"

### Kingdom-Wide CRISPR Guide Design with ALLEGRO (Supplementary Material)

All references in these supplementary materials are provided at the end of this document.

#### 1 Supplementary Notes

##### 1.1 Problem Definition

In the set covering problem, given a set $U$ of $n$ elements, and a set $S$ of $m$ subsets of $U$, the objective is to find the smallest collection of subsets whose union includes every element in $U$. The set covering problem can be formulated as an integer linear program (ILP), specifically a binary linear program, as it involves binary decisions on whether to include or exclude the subsets in the final collection. Solving ILPs is known to be NP-hard [1]. An approach to approximate the solution to an ILP is to relax its integer-constrained variables and allow them to take on fractional values. This transforms the ILP into a linear program (LP), which is solvable in polynomial time using algorithms such as the interior-point method [2] or the ellipsoid algorithm [3]. In this section, we (i) provide a mathematical formulation for our problem, (ii) describe the associated ILP, (iii) formulate the relaxation of ILP into the corresponding proxy LP (which is solvable in polynomial time), and (iv) describe a heuristic that allows us to discard redundant subsets to improve time and memory performance.

The inputs are (1) a set $U$ of species and (2) a set $G$ containing the DNA primary sequences of the genes of interest. While one would prefer to have the same set $G$ of orthologous genes in the species in $U$, it is likely that some species will be missing some genes (e.g., because of incomplete genome assemblies). We define any substring of length 20 in the set of strings $G$ which follows the protospacer adjacent motif (PAM) $\{A|C|G|T\}GG$ as a *guide*. We say that a guide $x$ targets species $u \in U$ if $x$ matches exactly the sequence of any of the genes in species $u$. Since the sequence of orthologous genes are similar (but not identical) across species, one can expect to find guides that can target the same gene in multiple species. The general problem we are interested in solving is the following. Given the inputs $U$ and $G$, find the smallest set of guides that target all species in $U$. This set of guides can be designed under certain constraints, which we call *tracks*. We defined two tracks, namely $A$ for *any* and $E$ for *each*. A track is also further characterized by the *multiplicity* factor $m$. In track $A_m$, the objective is to find the smallest set of guides that target every species at least $m$ times in any of the genes. For instance, if $|G| = 5$, the set of guides for track $A_6$ will have to target each species at least six times in any of the five genes. In track $A_m$, there are no restrictions on how many times each gene is targeted, and no guarantee that every gene will be hit at least once. The only requirement is that the total number of hits must be at least $m$ for each species. In track $E_m$, the objective is to find the smallest set of guides that target every gene in each species at least $m$ times. For instance, if $|G| = 5$, the set of guides for track $E_6$ will have to target each of the five genes at least six times (which is a total of 30 hits per species), in all species. By design, each guide in track $A_m$ may only count once per species in fulfilling the multiplicity requirement, regardless of how many times a guide targets the same species. In track $E_m$, a guide may only count once per gene toward the multiplicity regardless of how many times it may appear in the same gene.

#### 1.2 Linear Program for Track *A₁*

We use a variable $x_i$ for each candidate guide in the gene set $G$ of all species. We assume that there are $k$ candidate guides throughout all species. Variables $x_i$ are binary in the ILP, i.e. $x_i \in \{0, 1\}$, where $x_i = 1$ indicates that guide $i$ should be included in the library. We relax this requirement in the LP and allow these variables to take fractional values, i.e. $x_i \in [0, 1]$, so that we can solve the LP in polynomial time. In track $A_m$, (i) the genes in every species must be targeted at least $m$ times by the guides in the library and (ii) the objective is to minimize the number of guides in the library. The linear program for track $A_m$ is as follows

$$\text{Minimize} \sum_{i=1}^{k} x_i$$

$$\text{Subject to} \sum_{i:x_i \in C_u}^{n} x_i \geq m \quad \forall u \in U$$

$$x_i \in [0, 1] \quad i = 1, \ldots, k$$

where $C_u$ is the set of guides that target anywhere in the gene set of species $u \in U$, and $n$ is the total number of guides in species $u$.

#### 1.3 Linear Program for Track *E*$_m$

This track shares the same objective function as in track $A_m$. However, we need to modify the constraints to reflect the requirement of targeting each gene at least $m$ times. The LP for track $E_m$ is as follows

$$\text{Minimize} \sum_{i=1}^{k} x_i$$

$$\text{Subject to} \sum_{i:x_i \in C_{g,u}}^{n} x_i \geq m \quad \forall g \in G \quad \forall u \in U$$

$$x_i \in [0, 1] \quad i = 1, \ldots, k$$

where $C_{g,u}$ refers to the set of guides that target gene $g \in G$ of species $u \in U$, and $n$ is the total number of guides in gene $g$ in species $u$.

#### 1.4 Linear Program Incorporating Guide Activity

So far we have only been concerned with the size of the library. However, not every guide has the same cutting activity due to sequence composition, secondary structure, and many other factors. Several studies have devised rules, machine learning methods, thermodynamic calculations, and other strategies to predict the cutting efficiency of sgRNAs (see, e.g., [4, 5, 6, 7, 8, 9]). To represent the cutting efficiency of each guide, we introduce a new objective function which includes a weight $w_i$ for each guide $i$ (higher is better). In the new LP, we want to (1) maximize the overall cutting efficiency of the library, but (2) also keep the number of guides at a minimum, as follows

$$\text{Maximize} \sum_{i=1}^{k} w_i x_i \tag{1}$$

$$\text{Minimize} \sum_{i=1}^{k} x_i \tag{2}$$

Objectives (1) and (2) highlight a trade-off. To harmonize these competing objectives within a multi-objective linear program, we introduce a new parameter $\beta$, to act as a proxy for the desired cardinality of the library (i.e., the number of guides in the library). Thus, we replace objective (2) with a constraint that forces the size of the library to be at most $\beta$. The LP in this case is as follows

$$\text{Maximize} \sum_{i=1}^{k} w_i x_i$$

$$\text{Subject to} \begin{cases} \sum_{i:x_i \in C_u}^{n} & x_i \geq m \quad \text{if track } A_m \\ \sum_{i:x_i \in C_{g,u}}^{n} & x_i \geq m \quad \text{if track } E_m \\ \sum_{i=1}^{k} & x_i \leq \beta \end{cases}$$

$$x_i \in [0,1] \quad i = 1, \ldots, k$$

where $C_u$ is the set of guides that target anywhere in the gene set of species $u \in U$ and $C_{g,u}$ refers to the set of guides that target any gene $g \in G$ of species $u \in U$.

#### 1.5 Solving the (Integer) Linear Program

Depending on the selected track, multiplicity, and whether guide weights are incorporated, ALLEGRO sets up an LP with the appropriate objective function and constraints. The LP is solved using Google OR-Tools via a revised simplex algorithm [10, 11]. In the solution of the LP, each guide $i$ is assigned a fractional value $x_i \in [0, 1]$. Observe that if we obtain $x_i = 0$, guide $i$ should be excluded from the solution, while if $x_i = 1$, guide $i$ should definitely be included. The challenge arises when dealing with guides for which $0 < x_i < 1$, as their inclusion remains uncertain and requires further decision-making.

Randomized rounding is a common technique to address the issue of having non-integer variables from the LP. In randomized rounding, one can treat $x_i$ as the probability of including guide $i$ in the solution. We tested randomized rounding extensively by carrying out thousands of randomized trials, but the results were not satisfactory. Instead, we create a reduced ILP only for the guides $i$ for which $0 < x_i < 1$. The number of these fractional variables is often several orders of magnitudes smaller than the initial set of candidates used in the LP. Although finding the optimal solution to an ILP is NP-hard, this new program is sufficiently small that the heuristics available in [11, 12] produce a satisfactory solution. A detailed workflow is illustrated in **Supplementary Figure 1**.

#### 1.6 Discarding Redundant Guides

When running experiments on the entire transcriptome of thousands of species, ALLEGRO has to process and evaluate billions of potential guides. Generating and solving linear programs with billions of variables is computationally demanding not only in terms of time but other computational resources, such as main memory (RAM). However, many guides can be safely ignored prior to the linear formulation step. For instance, when running ALLEGRO for an $A_1$ experiment, if there is a set of guides that target exclusively a single species, we can keep only the best scoring guide in the set and discard the others.

To discard redundant guides without affecting the quality of the solution of the linear program, ALLEGRO uses a *redundancy threshold* $r$ that works as follows. For track $A_m$, if there is a set of guides $G$ that target exclusively the same set $S$ of species where $|S| \leq r$, only the top-$m$ scoring guides in $G$ are kept. For track $E_m$, if there is a set of guides $G$ that target exclusively the same set $S$ of genes where $|S| \leq r$, only the top-$m$ scoring guides in $G$ are kept. This technique significantly reduces the main memory requirement in large-scale experiments (e.g., our transcriptome-wide analysis for 1,000 species).

#### 2 Supplementary Figures

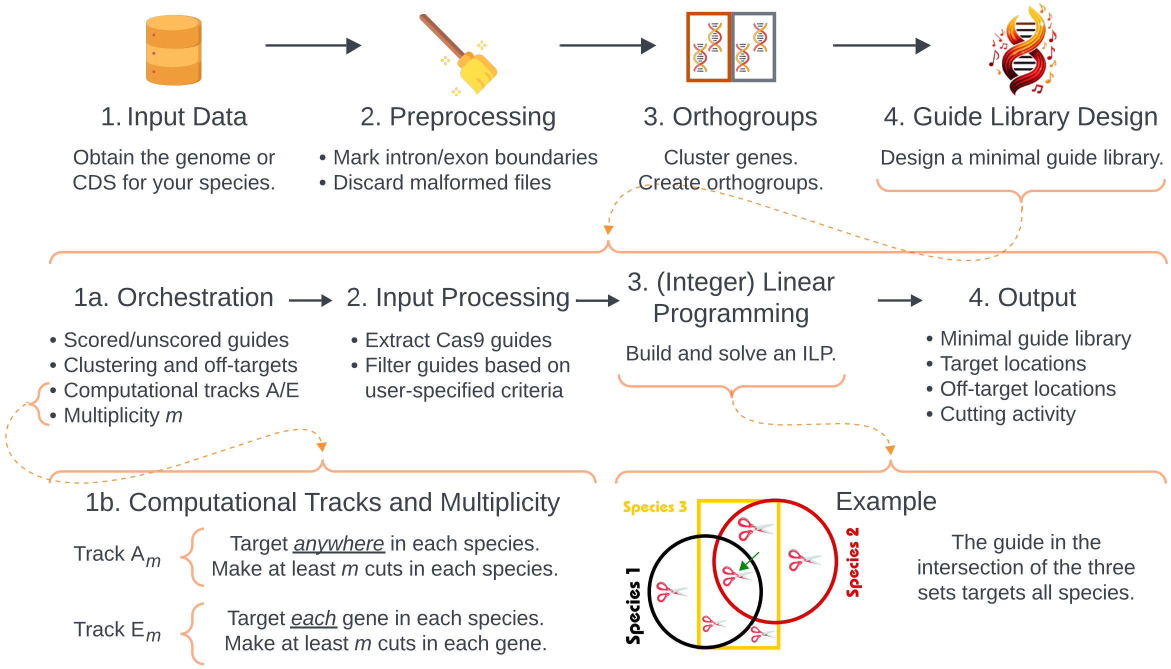


**Supplementary Figure 1**. Detailed overview of ALLEGRO. Once species are downloaded and preprocessed, they are used as input to ALLEGRO which designs a custom guide library based on a variety of user-defined parameters using linear programming.

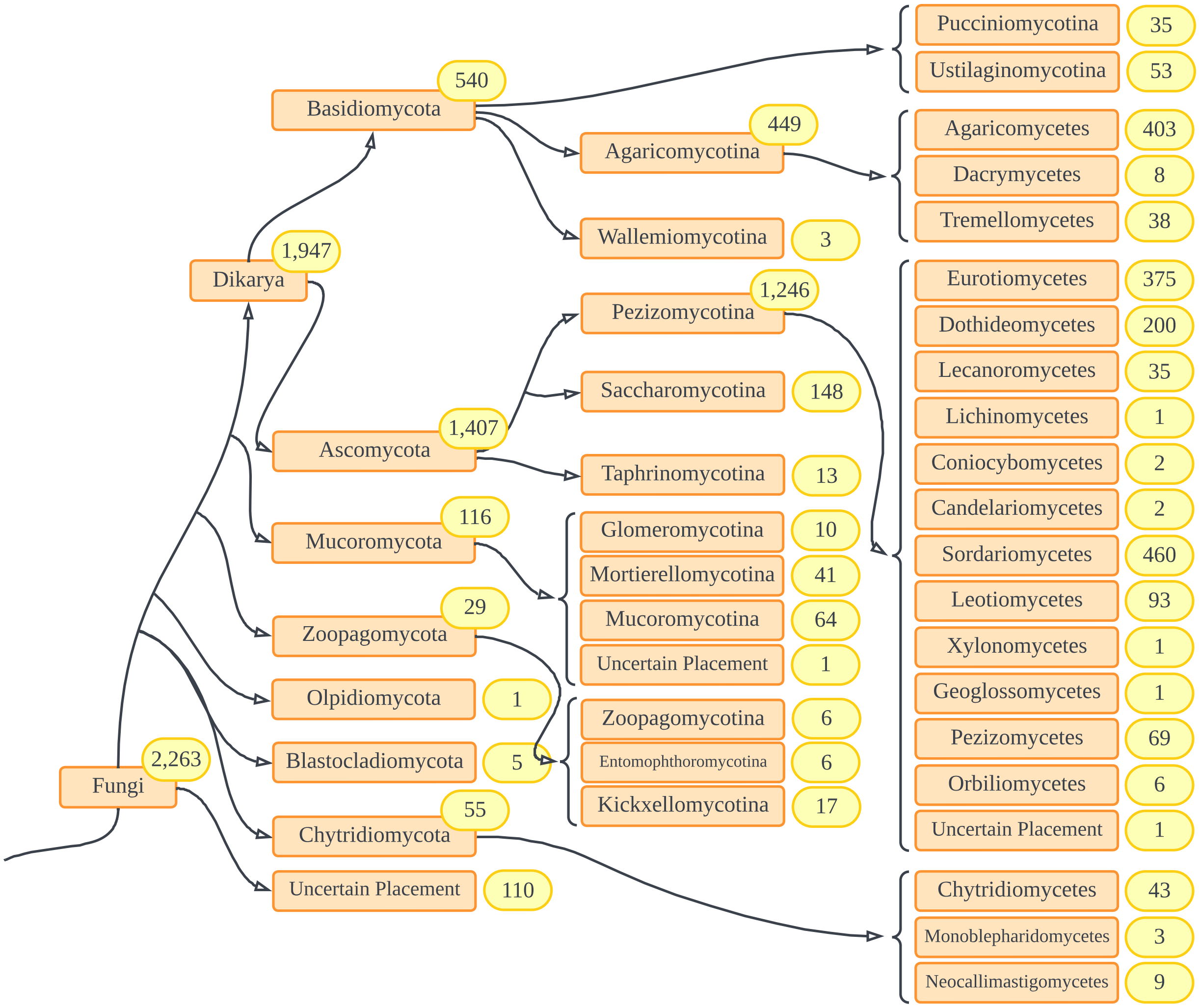


**Supplementary Figure 2**. A tree depicting the groupings of lineages for species used as input to ALLEGRO for designing sgRNAs targeting the six auxotrophic genes of interest. Numbers within the yellow oval boxes indicate the number of species in each group included in the input dataset. We created this tree based on [13, 14], and queried the NCBI Taxonomy database for species that were still unplaced [15]. We placed a species in the ”Uncertain Placement” group if none of the previous methods could identify its placement.

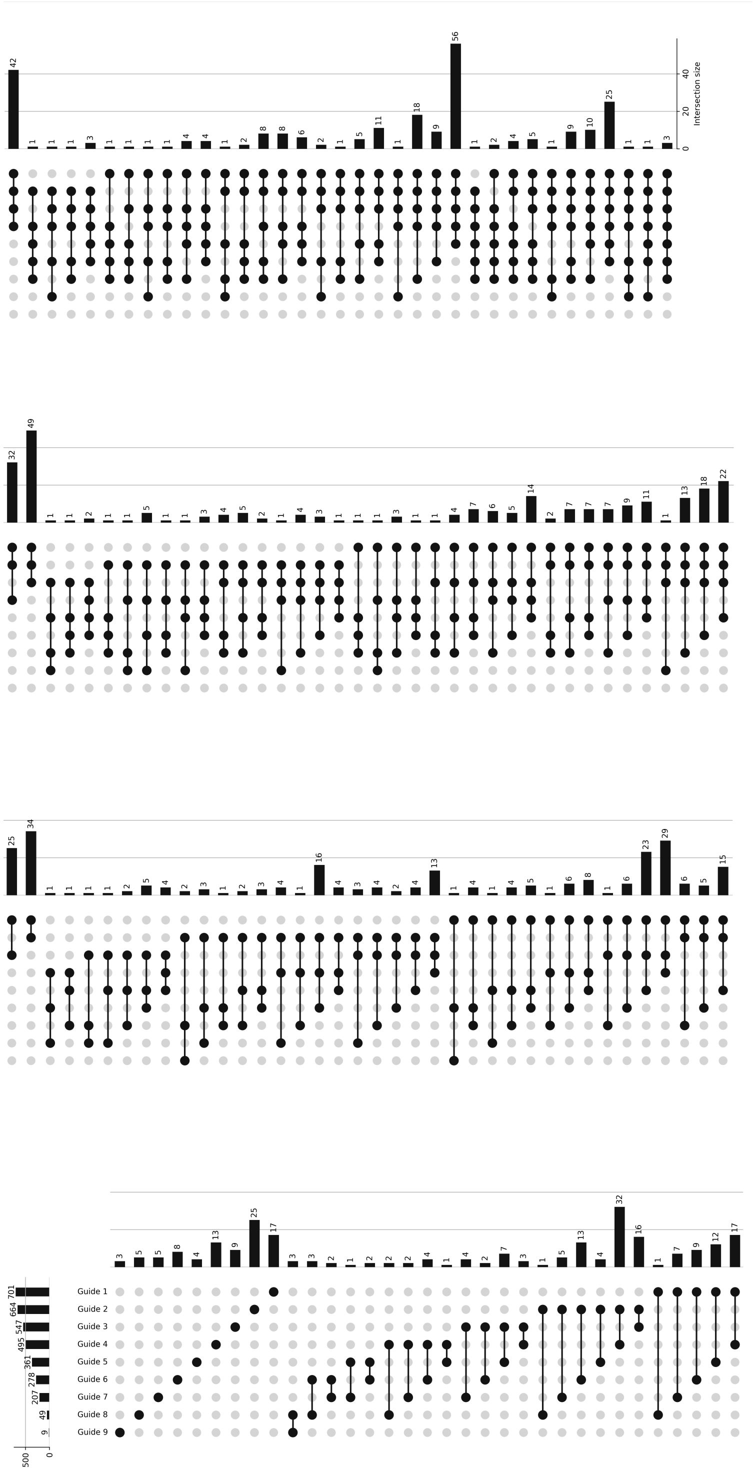


**Supplementary Figure 3**. Upset plot of the nine gRNAs that target anywhere in 1,000 Ascomycota species to fulfill track $A_1$, including the number of unique and overlapping target subsets. Created using the UpSet-Plot Python package [16].

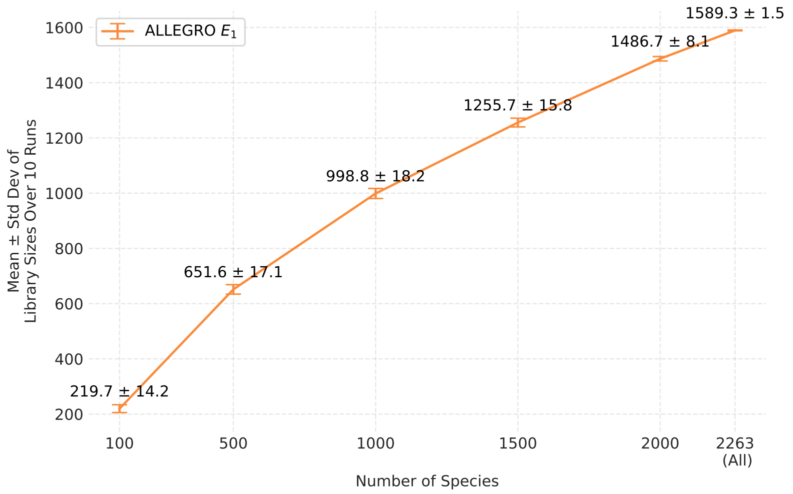


| X Values | Y Mean | Sample Std Dev (ddof=1) | CV (%) |
|---|---|---|---|
| 100 | 219.7 | 14.19741761 | 6.462183708 |
| 500 | 651.6 | 17.12827422 | 2.628648591 |
| 1000 | 998.8 | 18.18302016 | 1.8204866 |
| 1500 | 1255.7 | 15.80471097 | 1.258637491 |
| 2000 | 1486.7 | 8.056053624 | 0.541874865 |
| 2263 | 1589.3 | 1.494434118 | 0.094030964 |

**Supplementary Figure 4**. For each choice of species $x = \{100, 500, 1000, 1500, 2263\}$, we created ten samples of $x$ species to reduce the sampling bias for track $E_1$. Although ALLEGRO produces variable guide libraries in each run, the results are stable with coefficients of variation of under 10% (CV (%)).

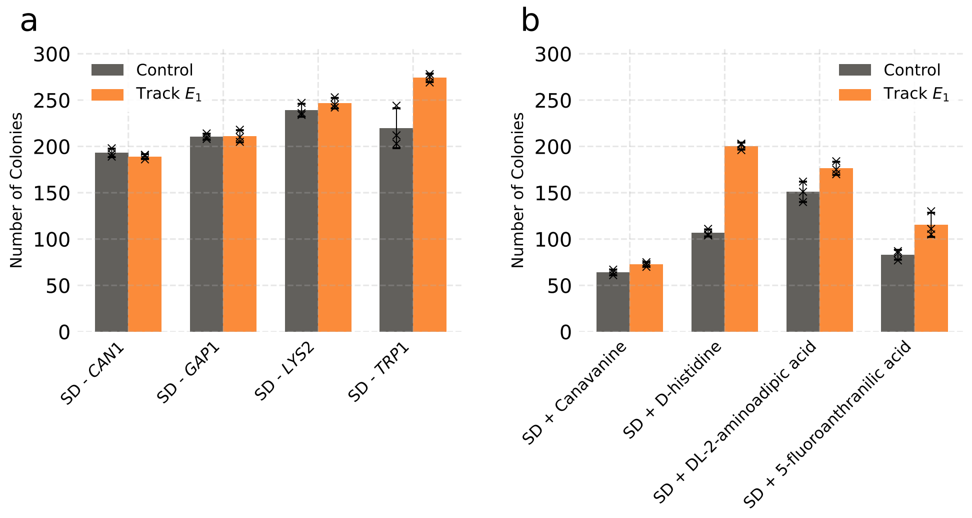


**Supplementary Figure 5**. *Y. lipolitica*'s number of transformants on -inhibitor (SD plates) and and +inhibitor conditions. **(a)** Number of colonies on -inhibitor plates for *Y. lipolytica* transformed with a plasmid expressing Cas9 and sgRNA from track $E_1$, targeting *CAN1*, *GAP1*, *LYS2*, and *TRP1* genes. **(b)** Number of colonies on +inhibitor plates for the same transformed colonies.

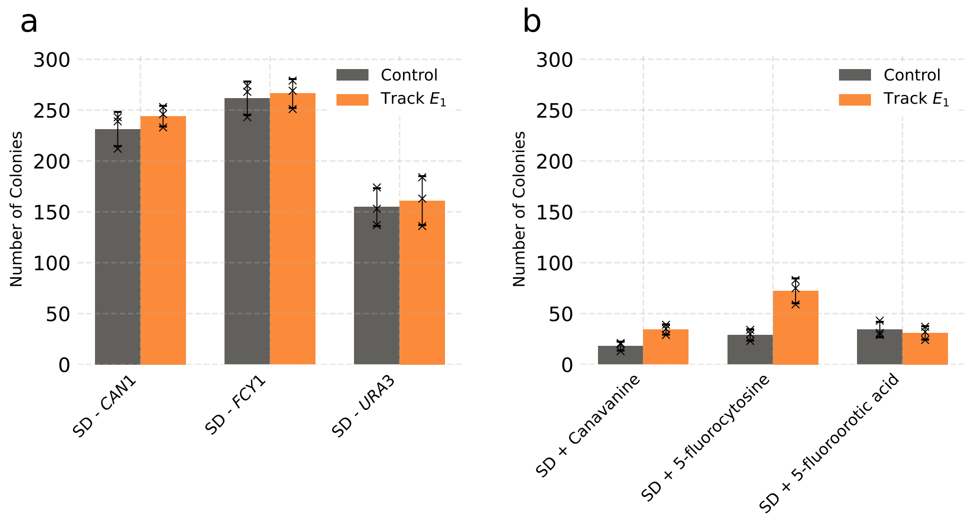


**Supplementary Figure 6**. *K. marxianus*'s number of transformants on -inhibitor (SD plates) and +inhibitor conditions. **(a)** Number of colonies on -inhibitor plates for *K. marxianus* transformed with plasmid expressing Cas9 and sgRNA from track $E_1$ targeting *CAN1* and *FCY1* genes, or with Cas9-RNP targeting *URA3* gene. **(b)** Number of colonies on +inhibitor plates for the same transformed colonies.

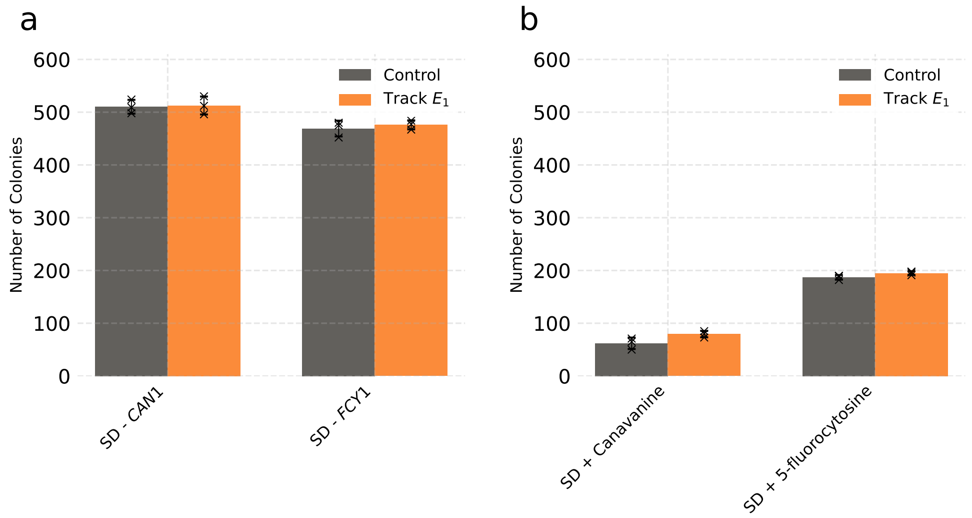


**Supplementary Figure 7**. *K. phaffii*'s number of transformants on -inhibitor (SD plates) and +inhibitor conditions. **(a)** Number of colonies on -inhibitor plates for *K. phaffii* transformed with a plasmid expressing Cas9 and sgRNA from track $E_1$, targeting *CAN1* and *FCY1* genes. **(b)** Number of colonies on +inhibitor plates for the same transformed colonies.

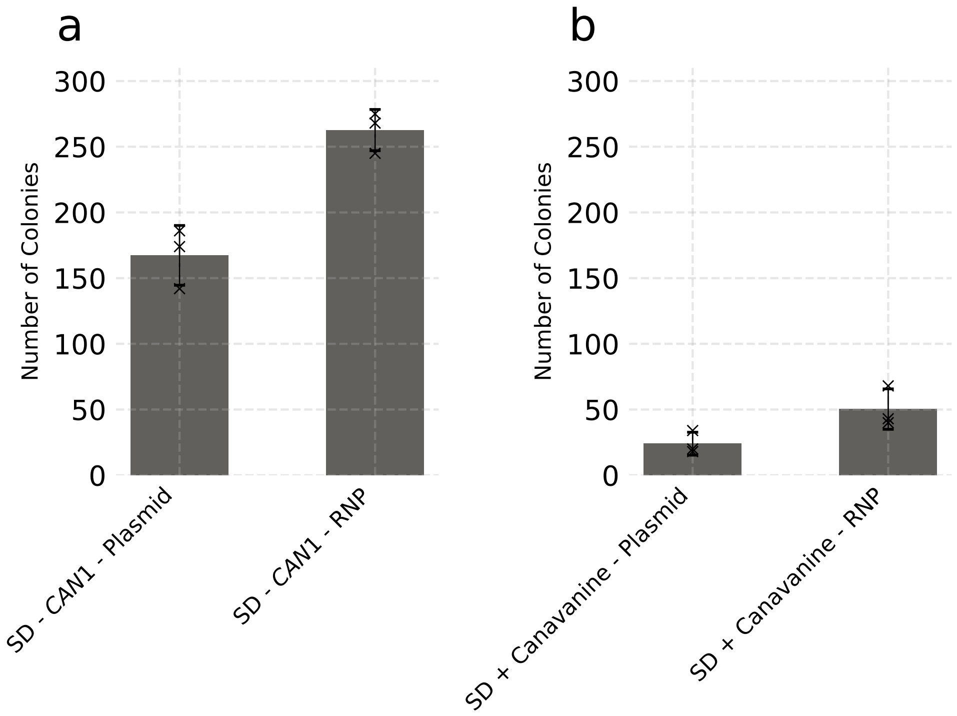


**Supplementary Figure 8**. Number of transformants under -inhibitor and +inhibitor conditions for *K. marxianus* protoplast transformation. **(a)** Number of colonies on -inhibitor plates for *K. marxianus* protoplasts transformed with both plasmid and Cas9-RNP complex containing sgRNA from track $E_1$, targeting the *CAN1* gene. **(b)** Number of colonies on +inhibitor plates for *K. marxianus* protoplasts transformed with plasmid and Cas9-RNP complex targeting the *CAN1* gene.

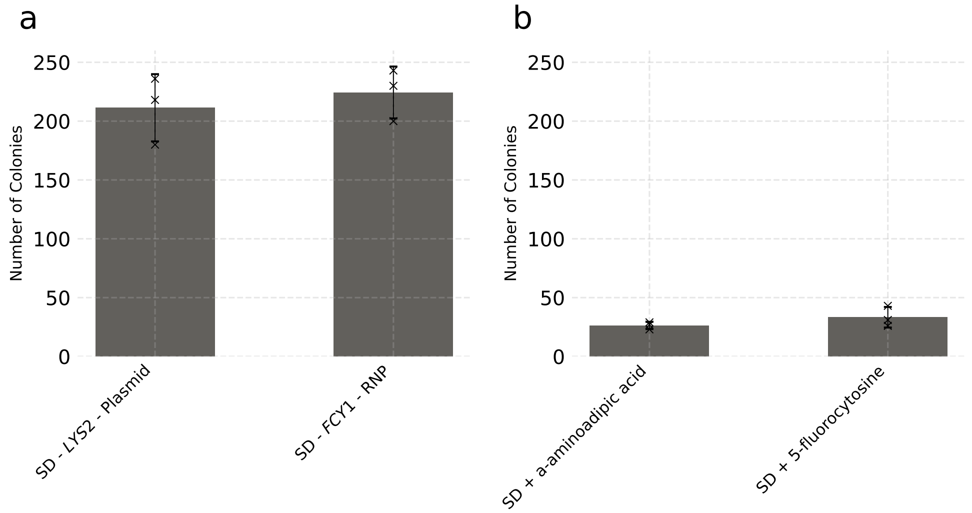


**Supplementary Figure 9**. Number of transformants under -inhibitor and +inhibitor conditions for *R. araucariae* transformation. **(a)** Number of colonies on -inhibitor plates for *R. araucariae* protoplasts transformed with Cas9-RNP complex containing sgRNA from track $E_1$, targeting the *FCY1* and *LYS2* genes. **(b)** Number of colonies on +inhibitor plates for *R. araucariae* protoplasts transformed with Cas9-RNP complex targeting the *FCY1* and *LYS2* genes.

### 3 Supplementary Tables

**Supplementary Table 1.** Strains used in this work.

| Strain | Description | Reference |
|---|---|---|
| *E. coli* TOP10 | F-mcrA$\Delta$ (mrr-hsdRMS-mcrBC)$\phi$80lacZ$\Delta$M15 $\Delta$lacX74 recA1 araD139 $\Delta$(*ara-leu*) 7697 galU galK rpsL (StrR) endA1 nupG $\lambda$ | Thermo Fisher Scientific |
| Ys302 | *K. marxianus* thermostable ATCC 26548 (CBS 6556) | ATCC |
| Ys402 | *K. marxianus* CBS6556 ($\Delta$*ura3*) | Lang et al. [17] |
| GS115 *his4::CAS9* | *K. phaffii* (*his4::CAS9*) | Tafrishi et al. [18] |
| Ys363 | *Y. lipolytica* PO1f (MatA, leu2-270, ura3-302, xpr2-322, axp-2) | ATCC |
| Y-17376 | *R. araucariae* | NRRL |

**Supplementary Table 2.** Plasmids used in this work.

| **Plasmid** | **Description** | **Reference** |
|---|---|---|
| pCRISPRpp | sgRNA library backbone, contains *PpHIS4* gene, *PARS1* and AmpR | Tafrishi et al. [18] |
| pSC012 | Easy clone *Y. lipolytica* Cas9 cutter without sgRNA contains *URA3* gene and AmpR | Robertson, et al. [19] |
| pIW601 | $P_{ScTEF1}$-KmCas9-SV40-ScCYC1t$P_{KmPRP1}$-tRNA$^{Gly}$-PspXI recognition site-*SUP4* | Wei S, et al. [20] |
| pRG03 | Ys402 *CAN1* KO track $E_1$-sgRNA in pIW601 | This work |
| pRG11 | Ys402 *FCY1* KO track $E_1$-sgRNA in pIW601 | This work |
| pRG21 | GS115 *CAN1* KO track $E_1$-sgRNA in pCRISPRpp | This work |
| pRG28 | GS115 *FCY1* KO track $E_1$-sgRNA in pCRISPRpp | This work |
| pRG36 | Ys363 *CAN1* KO track $E_1$-sgRNA in pSC012 | This work |
| pRG40 | Ys363 *TRP1* KO track $E_1$-sgRNA in pSC012 | This work |
| pRG46 | Ys363 *LYS2* KO track $E_1$-sgRNA in pSC012 | This work |
| pRG55 | Ys363 *GAP1* KO track $E_1$-sgRNA in pSC012 | This work |
| pRG01 | Ys402 *CAN1* KO control-sgRNA in pIW601 | This work |
| pRG10 | Ys402 *FCY1* KO control-sgRNA in pIW601 | This work |
| pRG20 | GS115 *CAN1* KO control-sgRNA in pCRISPRpp | This work |
| pRG26 | GS115 *FCY1* KO control-sgRNA in pCRISPRpp | This work |
| pRG34 | Ys363 *CAN1* KO control-sgRNA in pSC012 | This work |
| pRG38 | Ys363 *TRP1* KO control-sgRNA in pSC012 | This work |
| pRG44 | Ys363 *LYS2* KO control-sgRNA in pSC012 | This work |
| pRG53 | Ys363 *GAP1* KO control-sgRNA in pSC012 | This work |

**Supplementary Table 3 (Part 1/3).** Primers used in this work.

| Primer Name | Sequence | Description |
|---|---|---|
| K.m-C1-FOR | GGCACAGCCTTTCTTGGATTTTTG | Forward primer for *CAN1* knockouts, resulted by control sgRNA in *K. marxianus* |
| K.m-C1-REV | GTTCCCTGTGAAATGGTATGGTG | Reverse primer for *CAN1* knockouts, resulted by control sgRNA in *K. marxianus* |
| K.m-CE-FOR | GAGGAAGACACTGATATACGCGG | Forward primer for *CAN1* knockouts, resulted by track $E_1$ sgRNA in *K. marxianus* |
| K.m-CE-REV | GTGTCCAATAACGCACAAACTGT | Reverse primer for *CAN1* knockouts, resulted by track $E_1$ sgRNA in *K. marxianus* |
| K.m-F1-FOR | GCGATGGTCCTTATCCAGGATTA | Forward primer for *FCY1* knockouts, resulted by control sgRNA in *K. marxianus* |
| K.m-F1-REV | CCCCTTTAGTCTACCGGCATTTTC | Reverse primer for *FCY1* knockouts, resulted by control sgRNA in *K. marxianus* |
| K.m-FE-FOR | GGGAGGTGTTCCAATCGGTG | Forward primer for *FCY1* knockouts, resulted by track $E_1$ sgRNA in *K. marxianus* |
| K.m-FE-REV | CAGCGTTTGTCATCTACAACCTTC | Reverse primer for *FCY1* knockouts, resulted by track $E_1$ sgRNA in *K. marxianus* |
| K.m-U1-FOR | GTATACCGTATCGCCGAATGGT | Forward primer for *URA3* knockouts, resulted by control sgRNA in *K. marxianus*. |
| K.m-U1-REV | CCACAGTTCTGTATTGTTGTCCC | Reverse primer for *URA3* knockouts, resulted by control gRNA in *K. marxianus*. |
| K.m-UE-FOR | GTTACTCGGAAAGAGCAGCTG | Forward primer for *URA3* knockouts, resulted by track $E_1$ sgRNA in *K. marxianus*. |
| K.m-UE-REV | CAATGCCCGCACCAGTCACAC | Reverse primer for *URA3* knockouts, resulted by track $E_1$ sgRNA in *K. marxianus*. |
| K.p-C1-FOR | GAACACGCAGAAGGAGCCTG | Forward primer for *URA3* knockouts, resulted by control sgRNA in *K. phaffii*. |
| K.p-C1-REV | CGGCTTTAAAGATGAGACCTACGA | Reverse primer for *URA3* knockouts, resulted by control sgRNA in *K. phaffii*. |

**Supplementary Table 3, Continued (Part 2/3).** Primers used in this work.

| Primer Name | Sequence | Description |
|---|---|---|
| K.p-CE-FOR | GCCCTCGGAACTGTCTTTCTTG | Forward primer for *URA3* knockouts, resulted by track $E_1$ sgRNA in *K. phaffii*. |
| K.p-CE-REV | GTTCTGGTTGCAGCTAATTTCGTC | Reverse primer for *URA3* knockouts, resulted by track $E_1$ sgRNA in *K. phaffii*. |
| K.p-F1-FOR | GGCATTGACAACTTCAACACCC | Forward primer for *FCY1* knockouts, resulted by track $E_1$ and control sgRNA in *K. phaffii*. |
| K.p-F1-REV | GACATTCAGCGATGAAGACGG | Reverse primer for *FCY1* knockouts, resulted by track $E_1$ and control sgRNA in *K. phaffii*. |
| Y.l-C1-FOR | GAAAATTACAACACCAGTGGCCA | Forward primer for *CAN1* knockouts, resulted by control sgRNA in *Y. lipolytica*. |
| Y.l-C1-REV | GGCAAAGGAGCCGGTGAT | Reverse primer for *CAN1* knockouts, resulted by control sgRNA in *Y. lipolytica*. |
| K.p-F1-REV | GACATTCAGCGATGAAGACGG | Reverse primer for *FCY1* knockouts, resulted by track $E_1$ and control sgRNA in *K. phaffii*. |
| Y.l-C1-FOR | GAAAATTACAACACCAGTGGCCA | Forward primer for *CAN1* knockouts, resulted by control sgRNA in *Y. lipolytica*. |
| Y.l-C1-REV | GGCAAAGGAGCCGGTGAT | Reverse primer for *CAN1* knockouts, resulted by control sgRNA in *Y. lipolytica*. |
| Y.l-CE-FOR | GGAAAATTACAACACCAGTGGCCA | Forward primer for *CAN1* knockouts, resulted by track $E_1$ sgRNA in *Y. lipolytica*. |
| Y.l-CE-REV | GTAAAGACGGCAAAGGAGCC | Reverse primer for *CAN1* knockouts, resulted by track $E_1$ sgRNA in *Y. lipolytica*. |
| Y.l-T1-FOR | GACTCAACTCTCGCCACTGTC | Forward primer for *TRP1* knockouts, resulted by control sgRNA in *Y. lipolytica*. |
| Y.l-T1-REV | CCTGATCCAGTCGTCGGAT | Reverse primer for *TRP1* knockouts, resulted by control sgRNA in *Y. lipolytica*. |

**Supplementary Table 3, Continued (Part 3/3).** Primers used in this work.

| Primer Name | Sequence | Description |
|---|---|---|
| Y.l-TE-FOR | GACGTTGCACGGGAGATTG | Forward primer for *CAN1* knockouts, resulted by track $E_1$ sgRNA in *Y. lipolytica*. |
| Y.l-TE-REV | CCAGAAGCGAAAACGACTGG | Reverse primer for *CAN1* knockouts, resulted by track $E_1$ sgRNA in *Y. lipolytica*. |
| Y.l-L1-FOR | GACGTGATACCGAGACTCATGACC | Forward primer for *LYS2* knockouts, resulted by control sgRNA in *Y. lipolytica*. |
| Y.l-L1-REV | ATGGCGTAGAGAGCCAGAG | Reverse primer for *LYS2* knockouts, resulted by control sgRNA in *Y. lipolytica*. |
| Y.l-LE-FOR | CCAACTGGTTTGTGACTCCCG | Forward primer for *LYS2* knockouts, resulted by track $E_1$ sgRNA in *Y. lipolytica*. |
| Y.l-LE-REV | GCGTCAGTGTTCTGGACAAC | Reverse primer for *LYS2* knockouts, resulted by track $E_1$ sgRNA in *Y. lipolytica*. |
| Y.l-G1-FOR | GCCTACGGTATCATTGGATGC | Forward primer for *GAP1* knockouts, resulted by control sgRNA in *Y. lipolytica*. |
| Y.l-G1-REV | CCCTTGACTCCAAAGAGGTTGA | Reverse primer for *GAP1* knockouts, resulted by control sgRNA in *Y. lipolytica*. |
| Y.l-GE-FOR | GGTATCAACCAACTCTCCCCA | Forward primer for *GAP1* knockouts, resulted by track $E_1$ sgRNA in *Y. lipolytica*. |
| Y.l-GE-REV | CGATGAGAATGGCTGCCG | Reverse primer for *GAP1* knockouts, resulted by track $E_1$ sgRNA in *Y. lipolytica*. |
| R.a-LE-FOR | GCGCGAAAAGTTTGTCGACA | Forward primer for *LYS2* knockouts, resulted by track $E_1$ sgRNA in *R. araucariae*. |
| R.a-LE-REV | CTCGTCATCGCCCGAAAC | Reverse primer for *LYS2* knockouts, resulted by track $E_1$ sgRNA in *R. araucariae*. |
| R.a-FE-FOR | GAGGCGCTTTTCGCAACAC | Forward primer for *FCY1* knockouts, resulted by track $E_1$ sgRNA in *R. araucariae*. |
| R.a-FE-REV | CATTGCGTCCGCCTCTTAC | Reverse primer for *FCY1* knockouts, resulted by track $E_1$ sgRNA in *R. araucariae*. |

**Supplementary Table 4.** sgRNAs used in this work.

| sgRNA Sequence | Description | Target Genome - Gene |
|---|---|---|
| GGCATGAACTTGGCCTTGAA | Track $E_1$ sgRNA | Ys402 - *CAN1* |
| TCTCCATGTGATATGTGCAC | Track $E_1$ sgRNA | Ys402 - *FCY1* |
| GACAGGAAGTTTGCCGACAT | Track $E_1$ sgRNA | Ys302 - *URA3* |
| TTTAGATACTGGAGAAACCC | Track $E_1$ sgRNA | GS115 - *CAN1* |
| ACACATATGGCAAGGGCTCA | Track $E_1$ sgRNA | GS115 - *FCY1* |
| ATTGGTACCGGTCTCTTCAT | Track $E_1$ sgRNA | Ys363 - *CAN1* |
| CTCGACATTGTGCAGCTGCA | Track $E_1$ sgRNA | Ys363 - *TRP1* |
| GATGACCAGGTGAAGATTCG | Track $E_1$ sgRNA | Ys363 - *LYS2* |
| CATCTCCAGATGATTGCCAT | Track $E_1$ sgRNA | Ys363 - *GAP1* |
| GACGACCAGATCAAGATCCG | Track $E_1$ sgRNA | Y-17376 - *LYS2* |
| GAAGTGTGGAACGAGGACAT | Track $E_1$ sgRNA | Y-17376 - *FCY1* |
| TTGCCTGCACCACAAACCAT | Control sgRNA | Ys402 - *CAN1* |
| AAGGAGGGAGGTGTTCCAAT | Control sgRNA | Ys402 - *FCY1* |
| GCGCACGGTGAATACACTCG | Control sgRNA | Ys302 - *URA3* |
| ACTGGGCCTGCAATCTGTAA | Control sgRNA | GS115 - *CAN1* |
| GCAGCTCTAGTGTCCGAAGA | Control sgRNA | GS115 - *FCY1* |
| GGAGGAGCTCTCCAGCAGGC | Control sgRNA | Ys363 - *CAN1* |
| TCGACATGTCTACATCCCGT | Control sgRNA | Ys363 - *TRP1* |
| GTACTAATACAGTAACACAA | Control sgRNA | Ys363 - *LYS2* |
| CAGAATGGCAAACCACAGCA | Control sgRNA | Ys363 - *GAP1* |

**Supplementary Table 5.** Brief functional description of the auxotrophic genes, corresponding chemical inhibitors, and their applied concentrations in yeast screenings.

| **Target Gene** | **Description** | **Chemical Inhibitor** | **Concentration** | **Reference** |
|---|---|---|---|---|
| *CAN1* | Plasma membrane arginine permease. Mutation confers L-canavanine resistance. | L-canavanine sulfate salt | 50 or 60 mg/L | Yang, et al. [21] |
| *FCY1* | Cytosine deaminase, involved in the hydrolytic deamination of cytosine to uracil, and deamination of 5-fluorocytosine (5-FC) to 5-fluorouracil (5-FU). | 5-fluorocytosine (5-FC) | 1 mM | Smith, et al. [22] |
| *GAP1* | General amino acid permease, involved in uptake of amino acids and related compounds. | D-histidine | 10 mM | Regenberg, et al. [23] |
| *LYS2* | Encodes $\alpha$-aminoadipate reductase, crucial in the lysine biosynthesis pathway. | a-aminoadipic acid | 4.36 g/L | Keeney, et al. [24] |
| *TRP1* | Phosphoribosylanthranilate isomerase, involved in tryptophan biosynthesis. | 5-fluoroanthranilic acid (5-FAA) | 1.5 g/L | Cheon, et al. [25] |
| *URA3* | Orotidine-5'-phosphate decarboxylase, involved in pyrimidines synthesis. Mutation confers 5-fluoroorotic acid resistance. | 5-fluoroorotic Acid (5-FOA) | 1 g/L | Widlund, et al. [26] |
