## Supplementary Data 3 for "Kingdom-Wide CRISPR Guide Design with ALLEGRO"

**Total background genes: 4367**

| Guide | Num. Unique Targets (Foreground genes) | GO ID | GO Term | Occurence in Background | Occurence in Foreground | p-value (hypergeometric test) - Ascending ↓ |
|---|---|---|---|---|---|---|
| **ACTTCAGGAGCCATCCAATA** | 54 | **Ontology: Biological Process** | | | | |
| | | GO:0006468 | protein phosphorylation | 101 | 54 | 1.36E-96 |
| | | GO:0043408 | regulation of MAPK cascade | 37 | 34 | 1.42E-67 |
| | | GO:0023014 | signal transduction | 37 | 34 | 1.42E-67 |
| | | GO:0035556 | intracellular signal transduction | 55 | 34 | 1.42E-56 |
| | | 2 Others | N/A | N/A | N/A | > 0.05 |
| | | **Ontology: Molecular Function** | | | | |
| | | GO:0004672 | protein kinase activity | 101 | 54 | 1.36E-96 |
| | | GO:0004674 | protein serine/threonine kinase activity | 79 | 47 | 5.98E-82 |
| | | GO:0005524 | ATP binding | 260 | 54 | 2.56E-69 |
| | | GO:0022857 | transmembrane transporter activity | 20 | 1 | 2.21E-01 |
| | | **Ontology: Cellular Component** | | | | |
| | | GO:0005737 | cytoplasm | 404 | 47 | 2.00E-42 |
| | | GO:0016020 | membrane | 67 | 1 | 5.68E-01 |

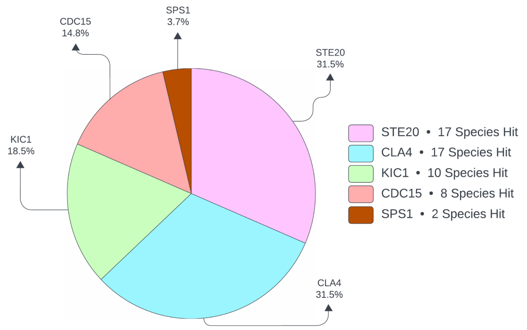


| Guide | Num. Unique Targets (Foreground genes) | GO ID | GO Term | Occurence in Background | Occurence in Foreground | p-value (hypergeometric test) - Ascending ↓ |
|---|---|---|---|---|---|---|
| **AGAGGAAGAGGAAGAGGAAG** | 1435 | **Ontology: Biological Process** | | | | |
| | | GO:0006281 | DNA repair | 38 | 32 | 7.80E-11 |
| | | GO:0016192 | vesicle-mediated transport | 32 | 27 | 2.33E-09 |
| | | GO:0006886 | intracellular protein transport | 30 | 25 | 1.57E-08 |
| | | GO:0000462 | maturation of SSU-rRNA from... | 26 | 22 | 7.04E-08 |
| | | GO:0006338 | chromatin remodeling | 24 | 20 | 4.76E-07 |
| | | GO:0006368 | transcription elongation by RNA... | 16 | 15 | 5.94E-07 |
| | | GO:0009267 | cellular response to starvation | 14 | 13 | 4.90E-06 |
| | | GO:0000723 | telomere maintenance | 13 | 12 | 1.40E-05 |
| | | GO:0016236 | macroautophagy | 13 | 12 | 1.40E-05 |
| | | GO:0034058 | endosomal vesicle fusion | 13 | 12 | 1.40E-05 |
| | | GO:0006122 | mitochondrial electron transport,... | 10 | 10 | 1.44E-05 |
| | | GO:0090630 | activation of GTPase activity | 9 | 9 | 4.39E-05 |
| | | GO:0030010 | establishment of cell polarity | 11 | 10 | 1.11E-04 |
| | | GO:0030490 | maturation of SSU-rRNA | 18 | 14 | 1.18E-04 |
| | | GO:0006406 | mRNA export from nucleus | 8 | 8 | 1.34E-04 |
| | | GO:0006325 | chromatin organization | 15 | 12 | 2.39E-04 |
| | | GO:0043044 | chromatin remodeling | 15 | 12 | 2.39E-04 |
| | | GO:0006623 | protein targeting to vacuole | 17 | 13 | 2.83E-04 |
| | | GO:0051382 | kinetochore assembly | 7 | 7 | 4.10E-04 |
| | | GO:0005975 | carbohydrate metabolic process | 16 | 12 | 6.70E-04 |
| | | GO:0042273 | ribosomal large subunit biogenesis | 18 | 13 | 7.13E-04 |
| | | GO:0007030 | Golgi organization | 9 | 8 | 8.56E-04 |
| | | GO:0006298 | mismatch repair | 9 | 8 | 8.56E-04 |
| | | GO:0051315 | attachment of mitotic spindle... | 6 | 6 | 1.25E-03 |
| | | GO:0044773 | mitotic DNA damage checkpoint signaling | 6 | 6 | 1.25E-03 |

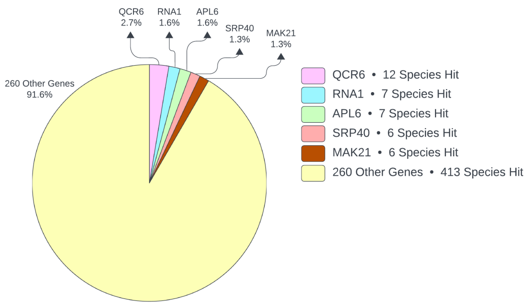

| | | | | |
|---|---|---|---|---|
| GO:0006359 | regulation of transcription by RNA... | 6 | 6 | 1.25E-03 |
| GO:0006364 | rRNA processing | 46 | 25 | 2.04E-03 |
| GO:0010833 | telomere maintenance via telomere... | 8 | 7 | 2.34E-03 |
| GO:0006891 | intra-Golgi vesicle-mediated transport | 8 | 7 | 2.34E-03 |
| GO:0000724 | double-strand break repair via... | 8 | 7 | 2.34E-03 |
| GO:0016567 | protein ubiquitination | 10 | 8 | 3.04E-03 |
| GO:0016226 | iron-sulfur cluster assembly | 10 | 8 | 3.04E-03 |
| GO:0006913 | nucleocytoplasmic transport | 12 | 9 | 3.41E-03 |
| GO:0006606 | protein import into nucleus | 12 | 9 | 3.41E-03 |
| GO:0001510 | RNA methylation | 14 | 10 | 3.54E-03 |
| GO:0043486 | obsolete histone exchange | 5 | 5 | 3.81E-03 |
| GO:0051455 | spindle attachment to meiosis I... | 5 | 5 | 3.81E-03 |
| GO:0042790 | nucleolar large rRNA transcription by... | 5 | 5 | 3.81E-03 |
| GO:0006361 | transcription initiation at RNA... | 5 | 5 | 3.81E-03 |
| GO:0031505 | fungal-type cell wall organization | 5 | 5 | 3.81E-03 |
| GO:0006449 | regulation of translational termination | 5 | 5 | 3.81E-03 |
| GO:0019511 | peptidyl-proline hydroxylation | 5 | 5 | 3.81E-03 |
| GO:0007094 | mitotic spindle assembly checkpoint... | 7 | 6 | 6.29E-03 |
| GO:0000463 | maturation of LSU-rRNA from... | 17 | 11 | 7.00E-03 |
| GO:0070475 | rRNA base methylation | 9 | 7 | 7.52E-03 |
| GO:0000398 | mRNA splicing, via spliceosome | 13 | 9 | 7.86E-03 |
| GO:0033314 | mitotic DNA replication checkpoint... | 4 | 4 | 1.16E-02 |
| GO:0045893 | positive regulation of DNA-templated... | 4 | 4 | 1.16E-02 |
| GO:0000479 | endonucleolytic cleavage of... | 4 | 4 | 1.16E-02 |
| GO:0007064 | mitotic sister chromatid cohesion | 4 | 4 | 1.16E-02 |
| GO:0031929 | TOR signaling | 4 | 4 | 1.16E-02 |
| GO:0072344 | rescue of stalled ribosome | 6 | 5 | 1.66E-02 |
| GO:0043328 | protein transport to vacuole involved... | 12 | 8 | 1.70E-02 |
| GO:0006260 | DNA replication | 12 | 8 | 1.70E-02 |
| GO:0000209 | protein polyubiquitination | 10 | 7 | 1.80E-02 |
| GO:0030833 | regulation of actin filament... | 3 | 3 | 3.54E-02 |
| GO:0000278 | mitotic cell cycle | 3 | 3 | 3.54E-02 |
| GO:0006384 | transcription initiation at RNA... | 3 | 3 | 3.54E-02 |
| GO:0000460 | maturation of 5.8S rRNA | 3 | 3 | 3.54E-02 |
| GO:0051754 | meiotic sister chromatid cohesion,... | 3 | 3 | 3.54E-02 |
| GO:1990758 | mitotic sister chromatid biorientation | 3 | 3 | 3.54E-02 |
| GO:0006265 | DNA topological change | 3 | 3 | 3.54E-02 |

| | | | | |
|---|---|---|---|---|
| GO:0000712 | resolution of meiotic recombination... | 3 | 3 | 3.54E-02 |
| GO:0032543 | mitochondrial translation | 3 | 3 | 3.54E-02 |
| GO:0007059 | chromosome segregation | 3 | 3 | 3.54E-02 |
| GO:0032790 | ribosome disassembly | 3 | 3 | 3.54E-02 |
| GO:0034501 | protein localization to kinetochore | 3 | 3 | 3.54E-02 |
| GO:0006272 | leading strand elongation | 3 | 3 | 3.54E-02 |
| GO:0031145 | anaphase-promoting complex-dependent... | 3 | 3 | 3.54E-02 |
| GO:0048478 | obsolete replication fork protection | 3 | 3 | 3.54E-02 |
| GO:0006888 | endoplasmic reticulum to Golgi... | 7 | 5 | 4.25E-02 |
| GO:0016573 | obsolete histone acetylation | 5 | 4 | 4.29E-02 |
| GO:1990116 | ribosome-associated... | 5 | 4 | 4.29E-02 |
| GO:0006335 | DNA replication-dependent chromatin... | 5 | 4 | 4.29E-02 |
| GO:0048208 | COPII vesicle coating | 5 | 4 | 4.29E-02 |
| GO:0006914 | autophagy | 5 | 4 | 4.29E-02 |
| GO:0070973 | protein localization to endoplasmic... | 5 | 4 | 4.29E-02 |
| GO:0031507 | heterochromatin formation | 5 | 4 | 4.29E-02 |
| GO:0006355 | regulation of DNA-templated... | 110 | 45 | 4.46E-02 |
| 306 Others | N/A | N/A | N/A | > 0.05 |
| **Ontology: Molecular Function** | | | | |
| GO:0005515 | protein binding | 207 | 135 | 1.60E-22 |
| GO:0042393 | histone binding | 23 | 19 | 1.22E-06 |
| GO:0031491 | nucleosome binding | 15 | 14 | 1.71E-06 |
| GO:0003682 | chromatin binding | 22 | 18 | 3.11E-06 |
| GO:0005519 | cytoskeletal regulatory protein binding | 9 | 9 | 4.39E-05 |
| GO:0140658 | ATP-dependent chromatin remodeler... | 13 | 11 | 1.80E-04 |
| GO:0008094 | ATP-dependent activity, acting on DNA | 10 | 9 | 3.10E-04 |
| GO:0140664 | ATP-dependent DNA damage sensor activity | 7 | 7 | 4.10E-04 |
| GO:0031267 | small GTPase binding | 12 | 10 | 4.69E-04 |
| GO:0005096 | GTPase activator activity | 14 | 11 | 5.90E-04 |
| GO:0016887 | ATP hydrolysis activity | 39 | 23 | 6.78E-04 |
| GO:0004402 | histone acetyltransferase activity | 11 | 9 | 1.20E-03 |
| GO:0034511 | U3 snoRNA binding | 6 | 6 | 1.25E-03 |
| GO:0016706 | 2-oxoglutarate-dependent dioxygenase... | 6 | 6 | 1.25E-03 |
| GO:0030983 | mismatched DNA binding | 6 | 6 | 1.25E-03 |
| GO:0017056 | structural constituent of nuclear pore | 13 | 10 | 1.43E-03 |
| GO:0003677 | DNA binding | 99 | 47 | 1.58E-03 |
| GO:0061630 | ubiquitin protein ligase activity | 14 | 10 | 3.54E-03 |
| GO:0001181 | RNA polymerase I general... | 5 | 5 | 3.81E-03 |

| | | | | |
|---|---|---|---|---|
| GO:0019237 | centromeric DNA binding | 5 | 5 | 3.81E-03 |
| GO:0031543 | peptidyl-proline dioxygenase activity | 5 | 5 | 3.81E-03 |
| GO:0000182 | rDNA binding | 5 | 5 | 3.81E-03 |
| GO:0031418 | L-ascorbic acid binding | 5 | 5 | 3.81E-03 |
| GO:0008173 | RNA methyltransferase activity | 13 | 9 | 7.86E-03 |
| GO:0008168 | methyltransferase activity | 20 | 12 | 1.13E-02 |
| GO:0003899 | DNA-directed 5'-3' RNA polymerase... | 4 | 4 | 1.16E-02 |
| GO:0001056 | RNA polymerase III activity | 4 | 4 | 1.16E-02 |
| GO:0030170 | pyridoxal phosphate binding | 4 | 4 | 1.16E-02 |
| GO:0001054 | RNA polymerase I activity | 4 | 4 | 1.16E-02 |
| GO:0000993 | RNA polymerase II complex binding | 4 | 4 | 1.16E-02 |
| GO:0003824 | catalytic activity | 25 | 14 | 1.41E-02 |
| GO:0046982 | protein heterodimerization activity | 16 | 10 | 1.42E-02 |
| GO:0016705 | oxidoreductase activity, acting on... | 6 | 5 | 1.66E-02 |
| GO:0003697 | single-stranded DNA binding | 6 | 5 | 1.66E-02 |
| GO:0042162 | telomeric DNA binding | 8 | 6 | 1.82E-02 |
| GO:0019887 | protein kinase regulator activity | 8 | 6 | 1.82E-02 |
| GO:0004386 | helicase activity | 8 | 6 | 1.82E-02 |
| GO:0022857 | transmembrane transporter activity | 20 | 11 | 3.36E-02 |
| GO:0016491 | oxidoreductase activity | 20 | 11 | 3.36E-02 |
| GO:0003906 | DNA-(apurinic or apyrimidinic site)... | 3 | 3 | 3.54E-02 |
| GO:0005507 | copper ion binding | 3 | 3 | 3.54E-02 |
| GO:0051015 | actin filament binding | 3 | 3 | 3.54E-02 |
| GO:0043022 | ribosome binding | 3 | 3 | 3.54E-02 |
| GO:0001164 | RNA polymerase I core promoter... | 3 | 3 | 3.54E-02 |
| GO:0047536 | 2-aminoadipate transaminase activity | 3 | 3 | 3.54E-02 |
| GO:0016791 | phosphatase activity | 9 | 6 | 3.94E-02 |
| GO:0003755 | peptidyl-prolyl cis-trans isomerase... | 7 | 5 | 4.25E-02 |
| GO:0016538 | cyclin-dependent protein... | 5 | 4 | 4.29E-02 |
| GO:0016747 | acyltransferase activity,... | 5 | 4 | 4.29E-02 |
| GO:0106388 | 18S rRNA... | 5 | 4 | 4.29E-02 |
| 267 Others | N/A | N/A | N/A | > 0.05 |
| **Ontology: Cellular Component** | | | | |
| GO:0030117 | membrane coat | 11 | 11 | 4.70E-06 |
| GO:0005770 | late endosome | 14 | 13 | 4.90E-06 |
| GO:0030897 | HOPS complex | 14 | 13 | 4.90E-06 |
| GO:0005750 | obsolete mitochondrial respiratory... | 10 | 10 | 1.44E-05 |
| GO:0051286 | cell tip | 9 | 9 | 4.39E-05 |
| GO:0000417 | HIR complex | 11 | 10 | 1.11E-04 |
| GO:0005759 | mitochondrial matrix | 7 | 7 | 4.10E-04 |
| GO:0030687 | preribosome, large subunit precursor | 18 | 13 | 7.13E-04 |
| GO:0048471 | perinuclear region of cytoplasm | 9 | 8 | 8.56E-04 |
| GO:0005956 | protein kinase CK2 complex | 6 | 6 | 1.25E-03 |

| | | | | |
|---|---|---|---|---|
| GO:0034456 | UTP-C complex | 6 | 6 | 1.25E-03 |
| GO:0016593 | Cdc73/Paf1 complex | 6 | 6 | 1.25E-03 |
| GO:0030690 | Noc1p-Noc2p complex | 6 | 6 | 1.25E-03 |
| GO:0030691 | Noc2p-Noc3p complex | 6 | 6 | 1.25E-03 |
| GO:0032300 | mismatch repair complex | 6 | 6 | 1.25E-03 |
| GO:0032389 | MutLalpha complex | 6 | 6 | 1.25E-03 |
| GO:0034399 | nuclear periphery | 6 | 6 | 1.25E-03 |
| GO:0005654 | nucleoplasm | 13 | 10 | 1.43E-03 |
| GO:0000776 | kinetochore | 8 | 7 | 2.34E-03 |
| GO:0031011 | Ino80 complex | 10 | 8 | 3.04E-03 |
| GO:0000785 | chromatin | 20 | 13 | 3.17E-03 |
| GO:0016586 | RSC-type complex | 12 | 9 | 3.41E-03 |
| GO:0005802 | trans-Golgi network | 5 | 5 | 3.81E-03 |
| GO:0031080 | nuclear pore outer ring | 5 | 5 | 3.81E-03 |
| GO:0080008 | Cul4-RING E3 ubiquitin ligase complex | 5 | 5 | 3.81E-03 |
| GO:0005739 | mitochondrion | 37 | 20 | 6.08E-03 |
| GO:0070187 | shelterin complex | 7 | 6 | 6.29E-03 |
| GO:0000781 | chromosome, telomeric region | 4 | 4 | 1.16E-02 |
| GO:0005622 | intracellular anatomical structure | 4 | 4 | 1.16E-02 |
| GO:0033255 | SAS acetyltransferase complex | 4 | 4 | 1.16E-02 |
| GO:0009986 | cell surface | 4 | 4 | 1.16E-02 |
| GO:0009277 | fungal-type cell wall | 4 | 4 | 1.16E-02 |
| GO:0005666 | RNA polymerase III complex | 4 | 4 | 1.16E-02 |
| GO:0005684 | U2-type spliceosomal complex | 4 | 4 | 1.16E-02 |
| GO:0043625 | delta DNA polymerase complex | 4 | 4 | 1.16E-02 |
| GO:0012507 | ER to Golgi transport vesicle membrane | 6 | 5 | 1.66E-02 |
| GO:0005643 | nuclear pore | 12 | 8 | 1.70E-02 |
| GO:0033565 | ESCRT-0 complex | 10 | 7 | 1.80E-02 |
| GO:0000329 | fungal-type vacuole membrane | 10 | 7 | 1.80E-02 |
| GO:0035267 | NuA4 histone acetyltransferase complex | 8 | 6 | 1.82E-02 |
| GO:0005886 | plasma membrane | 24 | 13 | 2.49E-02 |
| GO:0008622 | epsilon DNA polymerase complex | 3 | 3 | 3.54E-02 |
| GO:0008623 | CHRAC | 3 | 3 | 3.54E-02 |
| GO:0005618 | cell wall | 3 | 3 | 3.54E-02 |
| GO:0030123 | AP-3 adaptor complex | 3 | 3 | 3.54E-02 |
| GO:0033698 | Rpd3L complex | 3 | 3 | 3.54E-02 |
| GO:0030864 | cortical actin cytoskeleton | 3 | 3 | 3.54E-02 |
| GO:0005680 | anaphase-promoting complex | 3 | 3 | 3.54E-02 |
| GO:0031617 | kinetochore | 3 | 3 | 3.54E-02 |
| GO:0000500 | RNA polymerase I upstream activating... | 3 | 3 | 3.54E-02 |
| GO:0000127 | transcription factor TFIIIC complex | 3 | 3 | 3.54E-02 |
| GO:0005884 | actin filament | 3 | 3 | 3.54E-02 |
| GO:0071011 | precatalytic spliceosome | 3 | 3 | 3.54E-02 |
| GO:0000790 | chromatin | 11 | 7 | 3.56E-02 |
| GO:0000812 | Swr1 complex | 7 | 5 | 4.25E-02 |
| GO:0000151 | ubiquitin ligase complex | 7 | 5 | 4.25E-02 |

| | | | | |
|---|---|---|---|---|
| GO:0070971 | endoplasmic reticulum exit site | 5 | 4 | 4.29E-02 |
| GO:0017119 | Golgi transport complex | 5 | 4 | 4.29E-02 |
| GO:1990112 | RQC complex | 5 | 4 | 4.29E-02 |
| 155 Others | N/A | N/A | N/A | > 0.05 |

**CAACCCATTCCACCTGTATT** 9

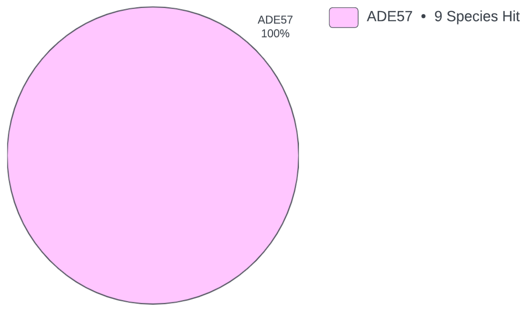


| | | | | |
|---|---|---|---|---|
| **Ontology: Biological Process** | | | | |
| GO:0046084 | adenine biosynthetic process | 9 | 9 | 6.33E-28 |
| GO:0009113 | purine nucleobase biosynthetic process | 9 | 9 | 6.33E-28 |
| GO:0006164 | purine nucleotide biosynthetic process | 9 | 9 | 6.33E-28 |
| GO:0006189 | de novo' IMP biosynthetic process | 9 | 9 | 6.33E-28 |
| **Ontology: Molecular Function** | | | | |
| GO:0004641 | phosphoribosylformyl glycinamidine... | 9 | 9 | 6.33E-28 |
| GO:0004637 | phosphoribosylamine -glycine ligase... | 9 | 9 | 6.33E-28 |
| GO:0046872 | metal ion binding | 21 | 9 | 1.86E-22 |
| GO:0005524 | ATP binding | 260 | 9 | 8.24E-12 |
| **Ontology: Cellular Component** | | | | |
| GO:0005829 | cytosol | 65 | 9 | 2.02E-17 |

**CCCACCTTCGACATTCCATT** 208

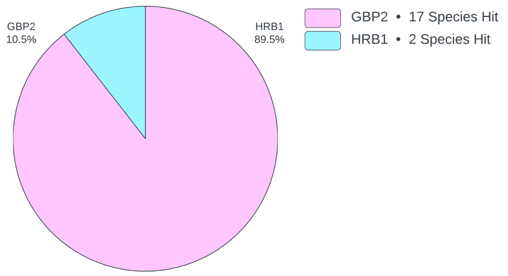


| | | | | |
|---|---|---|---|---|
| **Ontology: Biological Process** | | | | |
| GO:0006744 | ubiquinone biosynthetic process | 4 | 1 | 1.77E-01 |
| **Ontology: Molecular Function** | | | | |
| GO:0003729 | mRNA binding | 234 | 204 | 0.00E-02 |
| GO:0003676 | nucleic acid binding | 476 | 208 | 4.07E-222 |
| GO:0003723 | RNA binding | 872 | 208 | 9.00E-156 |
| GO:0008289 | lipid binding | 3 | 1 | 1.36E-01 |
| **Ontology: Cellular Component** | | | | |
| GO:0005737 | cytoplasm | 404 | 189 | 3.26E-191 |
| GO:0005634 | nucleus | 874 | 191 | 7.62E-119 |
| GO:1990904 | ribonucleoprotein complex | 212 | 116 | 8.49E-110 |

**GAAGGAGAAGAAGGAGAAGA** 908

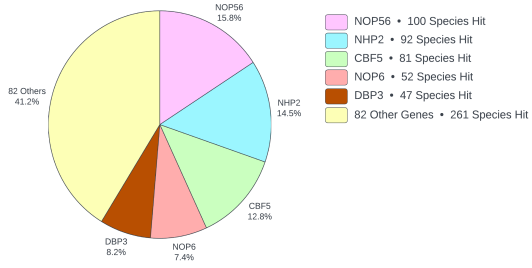


| | | | | |
|---|---|---|---|---|
| **Ontology: Biological Process** | | | | |
| GO:0031120 | snRNA pseudouridine synthesis | 82 | 82 | 5.78E-58 |
| GO:1990481 | mRNA pseudouridine synthesis | 82 | 82 | 5.78E-58 |
| GO:0031118 | rRNA pseudouridine synthesis | 82 | 82 | 5.78E-58 |
| GO:0009451 | RNA modification | 82 | 82 | 5.78E-58 |
| GO:0000495 | box H/ACA sno(s) RNA 3'-end processing | 82 | 82 | 5.78E-58 |
| GO:0042254 | ribosome biogenesis | 102 | 94 | 5.88E-56 |
| GO:0001522 | pseudouridine synthesis | 84 | 82 | 1.32E-54 |
| GO:0006396 | RNA processing | 90 | 84 | 3.77E-51 |
| GO:0042274 | ribosomal small subunit biogenesis | 37 | 36 | 4.74E-24 |
| GO:0000472 | endonucleolytic cleavage to generate... | 14 | 14 | 2.61E-10 |
| GO:0000480 | endonucleolytic cleavage in 5'-ETS of... | 14 | 14 | 2.61E-10 |
| GO:0000056 | ribosomal small subunit export from... | 16 | 14 | 2.04E-08 |
| GO:0006334 | nucleosome assembly | 19 | 15 | 8.83E-08 |
| GO:0045910 | negative regulation of DNA recombination | 10 | 10 | 1.45E-07 |
| GO:0030261 | chromosome condensation | 10 | 10 | 1.45E-07 |
| GO:0000447 | endonucleolytic cleavage in ITS1 to... | 20 | 15 | 2.86E-07 |

| | | | | |
|---|---|---|---|---|
| GO:0006360 | transcription by RNA polymerase I | 10 | 9 | 5.73E-06 |
| GO:0006364 | rRNA processing | 46 | 21 | 1.25E-04 |
| GO:0006418 | tRNA aminoacylation for protein... | 8 | 6 | 1.51E-03 |
| GO:0006417 | regulation of translation | 4 | 4 | 1.86E-03 |
| GO:0016579 | protein deubiquitination | 12 | 7 | 4.86E-03 |
| GO:1904802 | RITS complex assembly | 3 | 3 | 8.97E-03 |
| GO:0031048 | regulatory ncRNA-mediated... | 3 | 3 | 8.97E-03 |
| GO:0043039 | tRNA aminoacylation | 3 | 3 | 8.97E-03 |
| GO:0048015 | phosphatidylinositol-mediated signaling | 6 | 4 | 1.94E-02 |
| GO:0051209 | release of sequestered calcium ion... | 6 | 4 | 1.94E-02 |
| GO:0000470 | maturation of LSU-rRNA | 7 | 4 | 3.80E-02 |
| GO:0006629 | lipid metabolic process | 10 | 5 | 3.81E-02 |
| GO:0006464 | protein modification process | 2 | 2 | 4.32E-02 |
| GO:0030970 | retrograde protein transport, ER to... | 2 | 2 | 4.32E-02 |
| GO:0032007 | negative regulation of TOR signaling | 2 | 2 | 4.32E-02 |
| GO:0034976 | response to endoplasmic reticulum stress | 2 | 2 | 4.32E-02 |
| GO:0006435 | threonyl-tRNA aminoacylation | 2 | 2 | 4.32E-02 |
| GO:0051726 | regulation of cell cycle | 2 | 2 | 4.32E-02 |
| GO:0036503 | ERAD pathway | 2 | 2 | 4.32E-02 |
| 121 Others | N/A | N/A | N/A | > 0.05 |
| **Ontology: Molecular Function** | | | | |
| GO:0030515 | snoRNA binding | 143 | 142 | 1.44E-99 |
| GO:0009982 | pseudouridine synthase activity | 82 | 82 | 5.78E-58 |
| GO:0019843 | rRNA binding | 42 | 37 | 9.17E-21 |
| GO:0000049 | tRNA binding | 28 | 28 | 5.71E-20 |
| GO:0003676 | nucleic acid binding | 476 | 180 | 7.70E-20 |
| GO:0004831 | tyrosine-tRNA ligase activity | 27 | 27 | 2.81E-19 |
| GO:0030527 | structural constituent of chromatin | 16 | 14 | 2.04E-08 |
| GO:0031492 | nucleosomal DNA binding | 10 | 10 | 1.45E-07 |
| GO:0003690 | double-stranded DNA binding | 12 | 11 | 2.91E-07 |
| GO:0003723 | RNA binding | 872 | 236 | 3.91E-07 |
| GO:0003724 | RNA helicase activity | 36 | 19 | 2.10E-05 |
| GO:0140359 | ABC-type transporter activity | 10 | 7 | 1.09E-03 |
| GO:0005524 | ATP binding | 260 | 74 | 1.50E-03 |
| GO:0004812 | aminoacyl-tRNA ligase activity | 8 | 6 | 1.51E-03 |
| GO:0000166 | nucleotide binding | 11 | 7 | 2.46E-03 |
| GO:0042626 | ATPase-coupled transmembrane... | 9 | 6 | 3.74E-03 |
| GO:0004843 | cysteine-type deubiquitinase activity | 13 | 7 | 8.65E-03 |
| GO:0004198 | calcium-dependent cysteine-type... | 3 | 3 | 8.97E-03 |
| GO:0004435 | phosphatidylinositol phospholipase C... | 6 | 4 | 1.94E-02 |
| GO:0008081 | phosphoric diester hydrolase activity | 7 | 4 | 3.80E-02 |
| GO:0031386 | protein tag activity | 2 | 2 | 4.32E-02 |

| | | | | |
|---|---|---|---|---|
| GO:0004829 | threonine-tRNA ligase activity | 2 | 2 | 4.32E-02 |
| GO:0016433 | rRNA (adenine) methyltransferase... | 2 | 2 | 4.32E-02 |
| GO:0009383 | rRNA (cytosine-C5-)-methyltransferase... | 2 | 2 | 4.32E-02 |
| GO:0015035 | protein-disulfide reductase activity | 2 | 2 | 4.32E-02 |
| 93 Others | N/A | N/A | N/A | > 0.05 |
| **Ontology: Cellular Component** | | | | |
| GO:0032040 | small-subunit processome | 173 | 166 | 3.06E-108 |
| GO:0031428 | box C/D methylation guide snoRNP complex | 142 | 142 | 1.23E-101 |
| GO:0005730 | nucleolus | 260 | 186 | 1.19E-74 |
| GO:0031429 | box H/ACA snoRNP complex | 82 | 82 | 5.78E-58 |
| GO:0017102 | methionyl glutamyl tRNA synthetase... | 27 | 27 | 2.81E-19 |
| GO:1990904 | ribonucleoprotein complex | 212 | 94 | 1.90E-15 |
| GO:0000786 | nucleosome | 16 | 14 | 2.04E-08 |
| GO:0030688 | preribosome, small subunit precursor | 22 | 16 | 2.27E-07 |
| GO:0030686 | 90S preribosome | 31 | 18 | 5.83E-06 |
| GO:0005788 | endoplasmic reticulum lumen | 4 | 4 | 1.86E-03 |
| GO:0017101 | aminoacyl-tRNA synthetase multienzyme... | 2 | 2 | 4.32E-02 |
| GO:0033596 | TSC1-TSC2 complex | 2 | 2 | 4.32E-02 |
| GO:0022625 | cytosolic large ribosomal subunit | 2 | 2 | 4.32E-02 |
| 47 Others | N/A | N/A | N/A | > 0.05 |

**GACGGCTTCATGATCCACTC** 361

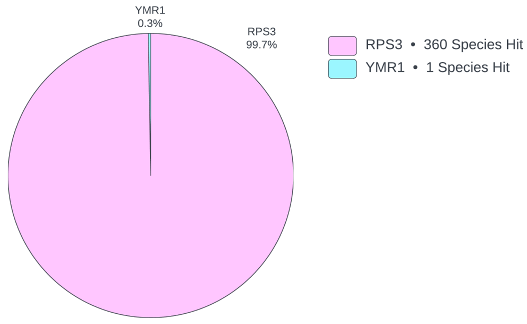


| | | | | |
|---|---|---|---|---|
| **Ontology: Biological Process** | | | | |
| GO:0006412 | translation | 1210 | 360 | 3.12E-218 |
| 2 Others | N/A | N/A | N/A | > 0.05 |
| **Ontology: Molecular Function** | | | | |
| GO:0003735 | structural constituent of ribosome | 363 | 360 | 0.00E+00 |
| GO:0003723 | RNA binding | 872 | 359 | 3.06E-278 |
| GO:0004438 | phosphatidylinositol-3-phosphate... | 1 | 1 | 8.27E-02 |
| **Ontology: Cellular Component** | | | | |
| GO:0022627 | cytosolic small ribosomal subunit | 359 | 359 | 0.00E+00 |
| GO:0015935 | small ribosomal subunit | 358 | 358 | 0.00E+00 |
| GO:0005634 | nucleus | 874 | 359 | 8.80E-278 |
| GO:0016020 | membrane | 67 | 1 | 9.97E-01 |

**GAGGCTGGTATCTCCAAGGA** 732

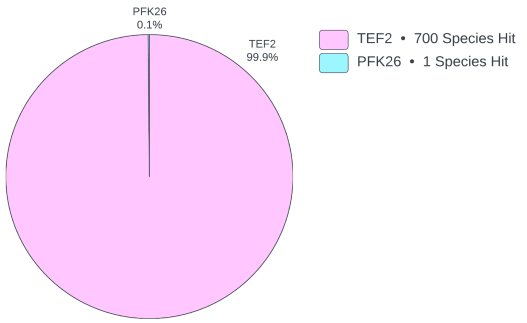


| | | | | |
|---|---|---|---|---|
| **Ontology: Biological Process** | | | | |
| GO:0006414 | translational elongation | 853 | 729 | 0.00E+00 |
| GO:0006412 | translation | 1210 | 728 | 0.00E+00 |
| 5 Others | N/A | N/A | N/A | > 0.05 |
| **Ontology: Molecular Function** | | | | |
| GO:0005525 | GTP binding | 868 | 730 | 0.00E+00 |
| GO:0003924 | GTPase activity | 865 | 730 | 0.00E+00 |
| GO:0003746 | translation elongation factor activity | 853 | 729 | 0.00E+00 |
| 10 Others | N/A | N/A | N/A | > 0.05 |
| **Ontology: Cellular Component** | | | | |
| 3 Others | N/A | N/A | N/A | > 0.05 |

**TCCAACTTCATCAAGAAGGT** 565

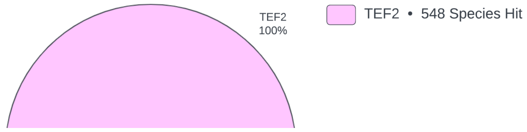


| | | | | |
|---|---|---|---|---|
| **Ontology: Biological Process** | | | | |
| GO:0006414 | translational elongation | 853 | 564 | 0.00E+00 |
| GO:0006412 | translation | 1210 | 563 | 0.00E+00 |
| 3 Others | N/A | N/A | N/A | > 0.05 |
| **Ontology: Molecular Function** | | | | |

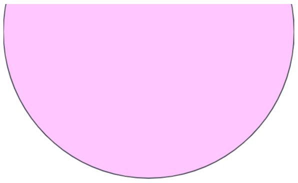

| | | | | |
|---|---|---|---|---|
| GO:0003924 | GTPase activity | 865 | 565 | 0.00E+00 |
| GO:0005525 | GTP binding | 868 | 564 | 0.00E+00 |
| GO:0003746 | translation elongation factor activity | 853 | 564 | 0.00E+00 |
| 3 Others | N/A | N/A | N/A | > 0.05 |
| **Ontology: Cellular Component** | | | | |
| N/A | N/A | N/A | N/A | nan |

**TGTTGTTGTTGTTGTTGTGG** 541

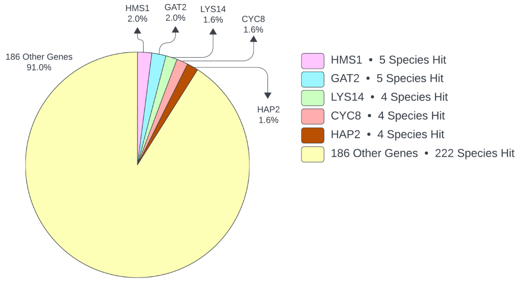


| | | | | |
|---|---|---|---|---|
| **Ontology: Biological Process** | | | | |
| GO:0006355 | regulation of DNA-templated... | 110 | 59 | 3.21E-26 |
| GO:0006357 | regulation of transcription by RNA... | 53 | 32 | 1.04E-16 |
| GO:0045944 | positive regulation of transcription... | 29 | 18 | 3.37E-10 |
| GO:0006351 | DNA-templated transcription | 35 | 16 | 1.05E-06 |
| GO:0006897 | endocytosis | 9 | 7 | 1.24E-05 |
| GO:0000290 | deadenylation-dependent decapping of... | 4 | 4 | 2.33E-04 |
| GO:0000184 | nuclear-transcribed mRNA catabolic... | 4 | 4 | 2.33E-04 |
| GO:0010468 | regulation of gene expression | 7 | 5 | 4.86E-04 |
| GO:0000122 | negative regulation of transcription... | 11 | 6 | 9.38E-04 |
| GO:0006487 | protein N-linked glycosylation | 5 | 4 | 1.05E-03 |
| GO:0032933 | SREBP signaling pathway | 3 | 3 | 1.89E-03 |
| GO:0098789 | obsolete pre-mRNA cleavage required... | 3 | 3 | 1.89E-03 |
| GO:0006031 | chitin biosynthetic process | 3 | 3 | 1.89E-03 |
| GO:0016197 | endosomal transport | 3 | 3 | 1.89E-03 |
| GO:0006038 | obsolete cell wall chitin... | 3 | 3 | 1.89E-03 |
| GO:0006654 | phosphatidic acid biosynthetic process | 6 | 4 | 2.85E-03 |
| GO:0033962 | P-body assembly | 7 | 4 | 5.99E-03 |
| GO:0018105 | peptidyl-serine phosphorylation | 4 | 3 | 6.87E-03 |
| GO:0030001 | metal ion transport | 4 | 3 | 6.87E-03 |
| GO:0044237 | cellular metabolic process | 2 | 2 | 1.53E-02 |
| GO:0006366 | transcription by RNA polymerase II | 2 | 2 | 1.53E-02 |
| GO:0007018 | microtubule-based movement | 2 | 2 | 1.53E-02 |
| GO:0030036 | actin cytoskeleton organization | 2 | 2 | 1.53E-02 |
| GO:0070059 | intrinsic apoptotic signaling pathway... | 2 | 2 | 1.53E-02 |
| GO:0010608 | post-transcriptional regulation of... | 2 | 2 | 1.53E-02 |
| GO:0000226 | microtubule cytoskeleton organization | 2 | 2 | 1.53E-02 |
| GO:0036498 | IRE1-mediated unfolded protein response | 2 | 2 | 1.53E-02 |
| GO:0001080 | nitrogen catabolite activation of... | 2 | 2 | 1.53E-02 |
| GO:0006772 | thiamine metabolic process | 2 | 2 | 1.53E-02 |
| GO:0044396 | actin cortical patch organization | 2 | 2 | 1.53E-02 |
| GO:0031124 | mRNA 3'-end processing | 2 | 2 | 1.53E-02 |
| GO:0007163 | establishment or maintenance of cell... | 2 | 2 | 1.53E-02 |
| GO:0000493 | box H/ACA snoRNP assembly | 2 | 2 | 1.53E-02 |

| | | | | |
|---|---|---|---|---|
| GO:0000294 | nuclear-transcribed mRNA catabolic... | 2 | 2 | 1.53E-02 |
| GO:0106035 | protein maturation by [4Fe-4S]... | 2 | 2 | 1.53E-02 |
| GO:0009229 | thiamine diphosphate biosynthetic... | 2 | 2 | 1.53E-02 |
| GO:0000147 | actin cortical patch assembly | 2 | 2 | 1.53E-02 |
| GO:0030968 | endoplasmic reticulum unfolded... | 5 | 3 | 1.56E-02 |
| GO:0007264 | small GTPase-mediated signal... | 5 | 3 | 1.56E-02 |
| GO:0007015 | actin filament organization | 5 | 3 | 1.56E-02 |
| GO:0006367 | transcription initiation at RNA... | 9 | 4 | 1.76E-02 |
| GO:0006511 | ubiquitin-dependent protein catabolic... | 10 | 4 | 2.65E-02 |
| GO:0055088 | lipid homeostasis | 6 | 3 | 2.83E-02 |
| GO:0007165 | signal transduction | 20 | 6 | 2.96E-02 |
| GO:0006468 | protein phosphorylation | 101 | 19 | 3.91E-02 |
| GO:0000387 | spliceosomal snRNP assembly | 3 | 2 | 4.22E-02 |
| GO:0006397 | mRNA processing | 3 | 2 | 4.22E-02 |
| GO:1904263 | positive regulation of TORC1 signaling | 3 | 2 | 4.22E-02 |
| GO:0010971 | positive regulation of G2/M... | 3 | 2 | 4.22E-02 |
| GO:0055085 | transmembrane transport | 39 | 9 | 4.48E-02 |
| GO:0015031 | protein transport | 7 | 3 | 4.51E-02 |
| 206 Others | N/A | N/A | N/A | > 0.05 |
| **Ontology: Molecular Function** | | | | |
| GO:0000981 | DNA-binding transcription factor... | 59 | 45 | 7.34E-30 |
| GO:0008270 | zinc ion binding | 83 | 41 | 8.98E-17 |
| GO:0000978 | RNA polymerase II cis-regulatory... | 30 | 22 | 1.69E-14 |
| GO:0043565 | sequence-specific DNA binding | 19 | 15 | 5.01E-11 |
| GO:0003700 | DNA-binding transcription factor... | 27 | 17 | 7.70E-10 |
| GO:0005515 | protein binding | 207 | 53 | 7.68E-08 |
| GO:0003677 | DNA binding | 99 | 32 | 1.20E-07 |
| GO:0008474 | palmitoyl-(protein) hydrolase activity | 7 | 7 | 4.33E-07 |
| GO:0035091 | phosphatidylinositol binding | 17 | 9 | 6.23E-05 |
| GO:0005509 | calcium ion binding | 8 | 6 | 7.91E-05 |
| GO:0003712 | transcription coregulator activity | 15 | 8 | 1.53E-04 |
| GO:0016787 | hydrolase activity | 19 | 9 | 1.88E-04 |
| GO:0003779 | actin binding | 9 | 6 | 2.13E-04 |
| GO:0043130 | ubiquitin binding | 16 | 8 | 2.73E-04 |
| GO:0000976 | transcription cis-regulatory region... | 14 | 7 | 6.75E-04 |
| GO:0046983 | protein dimerization activity | 11 | 6 | 9.38E-04 |
| GO:0032266 | phosphatidylinositol-3-phosphate binding | 5 | 4 | 1.05E-03 |
| GO:0051537 | 2 iron, 2 sulfur cluster binding | 5 | 4 | 1.05E-03 |
| GO:0052689 | carboxylic ester hydrolase activity | 3 | 3 | 1.89E-03 |
| GO:0051536 | iron-sulfur cluster binding | 3 | 3 | 1.89E-03 |
| GO:0042171 | lysophosphatidic acid acyltransferase... | 3 | 3 | 1.89E-03 |
| GO:0004100 | chitin synthase activity | 3 | 3 | 1.89E-03 |
| GO:0016251 | RNA polymerase II general... | 6 | 4 | 2.85E-03 |

| | | | | |
|---|---|---|---|---|
| GO:0030276 | clathrin binding | 4 | 3 | 6.87E-03 |
| GO:0003824 | catalytic activity | 25 | 8 | 8.31E-03 |
| GO:0050072 | obsolete m7G(5') pppN diphosphatase... | 2 | 2 | 1.53E-02 |
| GO:0051539 | 4 iron, 4 sulfur cluster binding | 2 | 2 | 1.53E-02 |
| GO:0140933 | 5'-(N(7)-methylguanosine... | 2 | 2 | 1.53E-02 |
| GO:0045504 | dynein heavy chain binding | 2 | 2 | 1.53E-02 |
| GO:0042134 | rRNA primary transcript binding | 2 | 2 | 1.53E-02 |
| GO:0030975 | thiamine binding | 2 | 2 | 1.53E-02 |
| GO:0004788 | thiamine diphosphokinase activity | 2 | 2 | 1.53E-02 |
| GO:0140662 | ATP-dependent protein folding chaperone | 2 | 2 | 1.53E-02 |
| GO:0004521 | RNA endonuclease activity | 2 | 2 | 1.53E-02 |
| GO:0016757 | glycosyltransferase activity | 5 | 3 | 1.56E-02 |
| GO:0019901 | protein kinase binding | 5 | 3 | 1.56E-02 |
| GO:0005543 | phospholipid binding | 5 | 3 | 1.56E-02 |
| GO:0016758 | hexosyltransferase activity | 5 | 3 | 1.56E-02 |
| GO:0031490 | chromatin DNA binding | 10 | 4 | 2.65E-02 |
| GO:0004674 | protein serine/threonine kinase activity | 79 | 16 | 3.03E-02 |
| GO:0046872 | metal ion binding | 21 | 6 | 3.73E-02 |
| GO:0004672 | protein kinase activity | 101 | 19 | 3.91E-02 |
| GO:0046873 | metal ion transmembrane transporter... | 3 | 2 | 4.22E-02 |
| GO:0008289 | lipid binding | 3 | 2 | 4.22E-02 |
| GO:0000009 | alpha-1,6-mannosyltransferase activity | 3 | 2 | 4.22E-02 |
| GO:0030145 | manganese ion binding | 3 | 2 | 4.22E-02 |
| GO:0000149 | SNARE binding | 3 | 2 | 4.22E-02 |
| GO:0051082 | unfolded protein binding | 3 | 2 | 4.22E-02 |
| 161 Others | N/A | N/A | N/A | > 0.05 |
| **Ontology: Cellular Component** | | | | |
| GO:0016020 | membrane | 67 | 23 | 2.33E-06 |
| GO:0017053 | transcription repressor complex | 4 | 4 | 2.33E-04 |
| GO:0043332 | mating projection tip | 4 | 4 | 2.33E-04 |
| GO:0016602 | CCAAT-binding factor complex | 4 | 4 | 2.33E-04 |
| GO:0000932 | P-body | 13 | 7 | 3.77E-04 |
| GO:0030428 | cell septum | 5 | 4 | 1.05E-03 |
| GO:0005847 | mRNA cleavage and polyadenylation... | 5 | 4 | 1.05E-03 |
| GO:0005739 | mitochondrion | 37 | 12 | 1.14E-03 |
| GO:0071944 | cell periphery | 3 | 3 | 1.89E-03 |
| GO:0016021 | membrane | 6 | 4 | 2.85E-03 |
| GO:0016592 | mediator complex | 6 | 4 | 2.85E-03 |
| GO:0005829 | cytosol | 65 | 16 | 4.70E-03 |
| GO:0005794 | Golgi apparatus | 7 | 4 | 5.99E-03 |
| GO:0030479 | actin cortical patch | 7 | 4 | 5.99E-03 |
| GO:0005669 | transcription factor TFIID complex | 7 | 4 | 5.99E-03 |
| GO:0005886 | plasma membrane | 24 | 8 | 6.32E-03 |
| GO:1990604 | IRE1-TRAF2-ASK1 complex | 2 | 2 | 1.53E-02 |

| | | | | |
|---|---|---|---|---|
| GO:0005868 | cytoplasmic dynein complex | 2 | 2 | 1.53E-02 |
| GO:0034967 | Set3 complex | 2 | 2 | 1.53E-02 |
| GO:0000172 | ribonuclease MRP complex | 2 | 2 | 1.53E-02 |
| GO:0000136 | mannan polymerase complex | 2 | 2 | 1.53E-02 |
| GO:0030125 | clathrin vesicle coat | 2 | 2 | 1.53E-02 |
| GO:0035974 | meiotic spindle pole body | 2 | 2 | 1.53E-02 |
| GO:0016514 | SWI/SNF complex | 2 | 2 | 1.53E-02 |
| GO:1990304 | MUB1-RAD6-UBR2 ubiquitin ligase complex | 2 | 2 | 1.53E-02 |
| GO:0000124 | SAGA complex | 10 | 4 | 2.65E-02 |
| GO:0005768 | endosome | 6 | 3 | 2.83E-02 |
| GO:0000808 | origin recognition complex | 3 | 2 | 4.22E-02 |
| GO:0070847 | core mediator complex | 3 | 2 | 4.22E-02 |
| 113 Others | N/A | N/A | N/A | > 0.05 |

**Total background genes: 4367**

| Guide | Num. Unique Targets (Foreground genes) | InterPro ID | GO Term | Occurence in Background | Occurence in Foreground | p-value (hypergeometric test) - Ascending ↓ |
|---|---|---|---|---|---|---|
| **ACTTCAGGAGCCATCCAATA** | 54 | **Protein Superfamily** | | | | |
| | | IPR011009 | Kinase-like_dom_sf | 109 | 54 | 4.25E-94 |
| | | **Protein Family** | | | | |
| | | IPR050285 | STE20_Ser/Thr_kinase | 37 | 34 | 1.42E-67 |
| | | IPR050629 | STE20/SPS1-PAK | 13 | 13 | 3.34E-26 |
| | | IPR050538 | MAP_kinase_kinase_kinase | 8 | 7 | 2.35E-13 |
| | | **Domain** | | | | |
| | | IPR000719 | Prot_kinase_dom | 96 | 54 | 2.64E-98 |
| | | IPR001245 | Ser-Thr/Tyr_kinase_cat_dom | 2 | 1 | 2.46E-02 |
| | | IPR020846 | MFS_dom | 11 | 1 | 1.28E-01 |
| | | **Other Features** | | | | |
| | | PIRSR000619-2 | PIRSR000619-2 | 64 | 52 | 2.63E-106 |
| | | PIRSR000628-2 | PIRSR000628-2 | 80 | 54 | 6.64E-105 |
| | | PIRSR500951-1 | PIRSR037393-1 | 73 | 53 | 1.59E-104 |
| | | PIRSR000620-1 | PIRSR000620-1 | 77 | 53 | 2.03E-102 |
| | | PIRSR000628-1 | PIRSR000628-1 | 80 | 53 | 5.69E-101 |
| | | PIRSR000620-2 | PIRSR000620-2 | 81 | 53 | 1.65E-100 |
| | | PIRSR500950-50 | PIRSR500950-50 | 76 | 52 | 3.00E-99 |
| | | PIRSR620777-50 | PIRSR620777-50 | 76 | 52 | 3.00E-99 |
| | | PIRSR000617-2 | PIRSR500947-50 | 64 | 50 | 5.90E-99 |
| | | PIRSR037393-1 | PIRSR000605-51 | 85 | 53 | 9.26E-99 |
| | | PIRSR000556-1 | PIRSR000556-1 | 85 | 53 | 9.26E-99 |
| | | PIRSR037921-1 | PIRSR037921-1 | 86 | 53 | 2.41E-98 |
| | | PIRSR000604-2 | PIRSR000604-2 | 78 | 52 | 2.77E-98 |
| | | PIRSR000636-2 | PIRSR000636-2 | 79 | 52 | 8.09E-98 |
| | | PIRSR000619-1 | PIRSR000619-1 | 79 | 52 | 8.09E-98 |
| | | PIRSR000666-1 | PIRSR000666-1 | 80 | 52 | 2.31E-97 |
| | | PIRSR000666-2 | PIRSR000666-2 | 73 | 51 | 2.92E-97 |
| | | PIRSR000552-1 | PIRSR000617-2 | 89 | 53 | 3.84E-97 |
| | | PIRSR630616-2 | PIRSR630616-2 | 90 | 53 | 9.33E-97 |
| | | PIRSR630616-1 | PIRSR037281-2 | 90 | 53 | 9.33E-97 |
| | | PIRSR500947-50 | PIRSR000556-2 | 74 | 51 | 9.39E-97 |
| | | PIRSR000604-1 | PIRSR000632-1 | 83 | 52 | 4.71E-96 |
| | | PIRSR500948-1 | PIRSR000559-2 | 76 | 51 | 8.91E-96 |
| | | PIRSR000631-1 | PIRSR000631-1 | 76 | 51 | 8.91E-96 |
| | | PIRSR000552-2 | PIRSR000552-2 | 78 | 51 | 7.61E-95 |
| | | PIRSR000605-50 | PIRSR000551-51 | 86 | 52 | 8.05E-95 |
| | | PIRSR038165-50 | PIRSR038165-50 | 86 | 52 | 8.05E-95 |
| | | PIRSR000559-1 | PIRSR000559-1 | 86 | 52 | 8.05E-95 |
| | | PIRSR000661-50 | PIRSR000661-50 | 86 | 52 | 8.05E-95 |
| | | PIRSR000550-3 | PIRSR000550-3 | 79 | 51 | 2.15E-94 |
| | | PIRSR000632-1 | PIRSR000660-1 | 79 | 51 | 2.15E-94 |
| | | PIRSR000550-1 | PIRSR000550-1 | 88 | 52 | 4.89E-94 |
| | | PIRSR630616-3 | PIRSR000632-2 | 88 | 52 | 4.89E-94 |
| | | PIRSR600239-51 | PIRSR600239-51 | 89 | 52 | 1.18E-93 |
| | | PIRSR628788-1 | PIRSR628788-1 | 90 | 52 | 2.78E-93 |
| | | G3DSA:1.10.510.10 | Transferase (Phosphotransferase) domain 1 | 101 | 53 | 6.55E-93 |
| | | PIRSR000632-2 | PIRSR038189-1 | 77 | 50 | 5.34E-92 |
| | | PIRSR037921-2 | PIRSR037921-2 | 77 | 50 | 5.34E-92 |
| | | PIRSR000554-1 | PIRSR000660-2 | 85 | 51 | 6.95E-92 |
| | | PIRSR000624-1 | PIRSR000624-1 | 71 | 49 | 1.31E-91 |
| | | PIRSR037568-1 | PIRSR630616-1 | 87 | 51 | 4.12E-91 |
| | | PIRSR000551-50 | PIRSR000551-50 | 88 | 51 | 9.80E-91 |

| | | | | | | |
|---|---|---|---|---|---|---|
| | | PIRSR000636-1 | PIRSR000636-1 | 80 | 50 | 1.08E-90 |
| | | PIRSR637770-1 | PIRSR637770-1 | 91 | 51 | 1.20E-89 |
| | | PIRSR000617-1 | PIRSR037281-3 | 83 | 50 | 1.81E-89 |
| | | PIRSR037014-2 | PIRSR037014-2 | 71 | 48 | 1.99E-88 |
| | | PIRSR038172-1 | PIRSR038172-1 | 88 | 50 | 1.41E-87 |
| | | PIRSR637770-2 | PIRSR630220-1 | 92 | 50 | 3.50E-86 |
| | | PIRSR628788-2 | PIRSR000605-50 | 85 | 49 | 1.29E-85 |
| | | PIRSR037014-1 | PIRSR037014-1 | 86 | 49 | 2.99E-85 |
| | | PIRSR037993-1 | PIRSR000624-2 | 86 | 49 | 2.99E-85 |
| | | PIRSR037393-2 | PIRSR037393-2 | 70 | 46 | 8.69E-83 |
| | | PIRSR000556-2 | PIRSR630616-3 | 71 | 46 | 2.46E-82 |
| | | PIRSR037993-2 | PIRSR500948-1 | 77 | 46 | 7.84E-80 |
| | | PIRSR038172-2 | PIRSR038172-2 | 64 | 44 | 1.01E-79 |
| | | PIRSR630220-1 | PIRSR628788-2 | 62 | 43 | 8.62E-78 |
| | | PIRSR000624-2 | PIRSR637770-2 | 63 | 43 | 2.71E-77 |
| | | PIRSR000554-2 | PIRSR000554-2 | 72 | 42 | 1.15E-70 |
| | | PIRSR630220-2 | PIRSR630220-2 | 54 | 39 | 1.86E-70 |
| | | PIRSR000551-51 | PIRSR500948-2 | 66 | 39 | 5.05E-65 |
| | | cd06614 | STKc_PAK | 37 | 32 | 3.20E-61 |
| | | PIRSR000660-1 | PIRSR037993-2 | 87 | 40 | 6.73E-61 |
| | | PIRSR037568-2 | PIRSR037568-2 | 63 | 36 | 1.64E-58 |
| | | PIRSR000661-51 | PIRSR000661-51 | 48 | 33 | 3.89E-57 |
| | | PIRSR000559-2 | disorder_prediction | 58 | 33 | 5.92E-53 |
| | | PIRSR000606-51 | PIRSR000605-50 | 73 | 34 | 1.15E-50 |
| | | PIRSR000605-51 | PIRSR000604-1 | 40 | 22 | 7.61E-34 |
| | | PIRSR038165-51 | PIRSR038165-51 | 11 | 11 | 3.51E-22 |
| | | cd06627 | STKc_Cdc7_like | 8 | 8 | 3.19E-16 |
| | | PIRSR620777-51 | PIRSR620777-51 | 11 | 7 | 9.41E-12 |
| | | PIRSR037281-3 | PIRSR000619-1 | 88 | 9 | 9.18E-07 |
| | | PIRSR000617-4 | PIRSR000617-4 | 7 | 3 | 6.04E-05 |
| | | PIRSF000654 | PIRSR000552-1 | 2 | 2 | 1.50E-04 |
| | | PIRSR037281-2 | PIRSR037281-2 | 76 | 4 | 1.39E-02 |
| | | PIRSR037281-1 | PIRSR037281-1 | 89 | 4 | 2.35E-02 |
| | | NON_CYTOPLASMIC_DOMAIN | PIRSR600239-51 | 504 | 1 | 9.99E-01 |
| **AGAGGAAGAGGAAGAGGAAG** | 1435 | **Protein Superfamily** | | | | |
| | | IPR036603 | RBP11-like | 4 | 4 | 1.16E-02 |
| | | 50 Others | N/A | N/A | N/A | > 0.05 |
| | | **Protein Family** | | | | |
| | | IPR003422 | Cyt_b-c1_6 | 10 | 10 | 1.44E-05 |
| | | IPR026739 | AP_beta | 9 | 9 | 4.39E-05 |
| | | IPR027038 | RanGap | 8 | 8 | 1.34E-04 |
| | | IPR040155 | CEBPZ/Mak21-like | 6 | 6 | 1.25E-03 |
| | | IPR040458 | Vid27 | 6 | 6 | 1.25E-03 |
| | | IPR000704 | Casein_kinase_II_reg-sub | 6 | 6 | 1.25E-03 |
| | | IPR045111 | Vps41/Vps8 | 13 | 10 | 1.43E-03 |
| | | IPR051842 | uS12_prolyl_hydroxylase | 5 | 5 | 3.81E-03 |
| | | IPR039754 | Esf1 | 5 | 5 | 3.81E-03 |
| | | IPR029523 | INO80B/Ies2 | 7 | 6 | 6.29E-03 |
| | | IPR039776 | Pds5 | 4 | 4 | 1.16E-02 |
| | | IPR051494 | BSD_domain-containing | 4 | 4 | 1.16E-02 |
| | | IPR039761 | Bms1/Tsr1 | 4 | 4 | 1.16E-02 |
| | | IPR031101 | Ctr9 | 4 | 4 | 1.16E-02 |
| | | IPR033194 | MFAP1 | 4 | 4 | 1.16E-02 |
| | | IPR005343 | Noc2 | 6 | 5 | 1.66E-02 |
| | | IPR008610 | Ebp2 | 6 | 5 | 1.66E-02 |
| | | IPR051377 | DNA_Pol-Epsilon_Subunit | 3 | 3 | 3.54E-02 |
| | | IPR039601 | Rrn5 | 3 | 3 | 3.54E-02 |
| | | IPR052414 | U3_snoRNA-assoc_WDR | 3 | 3 | 3.54E-02 |
| | | IPR009316 | COG2 | 3 | 3 | 3.54E-02 |
| | | IPR026740 | AP3_beta | 3 | 3 | 3.54E-02 |

| | | | | | | |
|---|---|---|---|---|---|---|
| | | IPR051526 | Beta-Glucosidase_SUN | 3 | 3 | 3.54E-02 |
| | | IPR052007 | Bud4 | 3 | 3 | 3.54E-02 |
| | | IPR051859 | DCAF | 3 | 3 | 3.54E-02 |
| | | IPR006958 | Mak16 | 3 | 3 | 3.54E-02 |
| | | IPR015661 | Bub1/Mad3 | 3 | 3 | 3.54E-02 |
| | | IPR037382 | Rsc/polybromo | 9 | 6 | 3.94E-02 |
| | | IPR022968 | Tsr3-like | 5 | 4 | 4.29E-02 |
| | | IPR053030 | Ribosomal_biogenesis_FAF1-like | 5 | 4 | 4.29E-02 |
| | | IPR028386 | CENP-C/Mif2/cnp3 | 5 | 4 | 4.29E-02 |
| | | IPR024880 | Sec16 | 5 | 4 | 4.29E-02 |
| | | 232 Others | N/A | N/A | N/A | > 0.05 |
| | | **Domain** | | | | |
| | | IPR019601 | Oxoglutarate/Fe-dep_Oase_C | 5 | 4 | 4.29E-02 |
| | | 38 Others | N/A | N/A | N/A | > 0.05 |
| | | **Domain** | | | | |
| | | mobidb-lite | disorder_prediction | 3393 | 1279 | 1.91E-40 |
| | | Coil | Coil | 2135 | 828 | 2.20E-16 |
| | | PTHR35711 | EXPRESSED PROTEIN | 17 | 14 | 3.76E-05 |
| | | PTHR45852 | SER/THR-PROTEIN KINASE RIO2 | 5 | 5 | 3.81E-03 |
| | | PTHR18934 | ATP-DEPENDENT RNA HELICASE | 4 | 4 | 1.16E-02 |
| | | PTHR44167 | PIRSR037993-1 | 6 | 5 | 1.66E-02 |
| | | PTHR23405 | MAINTENANCE OF KILLER 16 MAK16 PROTEIN-RELATED | 3 | 3 | 3.54E-02 |
| | | cd00609 | AAT_like | 3 | 3 | 3.54E-02 |
| | | PTHR43670 | HEAT SHOCK PROTEIN 26 | 3 | 3 | 3.54E-02 |
| | | 136 Others | N/A | N/A | N/A | > 0.05 |
| **CAACCCATTCCACCTGTATT** | 9 | **Protein Family** | | | | |
| | | IPR000115 | PRibGlycinamide_synth | 9 | 9 | 6.33E-28 |
| | | IPR004733 | PurM_cligase | 9 | 9 | 6.33E-28 |
| | | **Domain** | | | | |
| | | IPR020561 | PRibGlycinamid_synth_ATP-grasp | 9 | 9 | 6.33E-28 |
| | | IPR011761 | ATP-grasp | 10 | 9 | 6.33E-27 |
| | | **Other Features** | | | | |
| | | SM01209 | GARS_A_3 | 9 | 9 | 6.33E-28 |
| | | SSF56059 | Glutathione synthetase ATP-binding domain-like | 12 | 9 | 1.39E-25 |
| | | NON_CYTOPLASMIC_DOMAIN | Non cytoplasmic domain | 504 | 1 | 6.69E-01 |
| **CCCACCTTCGACATTCCATT** | 208 | **Protein Superfamily** | | | | |
| | | IPR012677 | Nucleotide-bd_a/b_plait_sf | 315 | 204 | 7.84E-262 |
| | | IPR035979 | RBD_domain_sf | 316 | 199 | 2.65E-246 |
| | | **Protein Family** | | | | |
| | | IPR050374 | RRT5_SRSF_SR | 191 | 147 | 9.47E-182 |
| | | **Domain** | | | | |
| | | IPR000504 | RRM_dom | 310 | 192 | 8.19E-230 |
| | | **Other Features** | | | | |
| | | cd00590 | RRM_SF | 47 | 42 | 5.36E-52 |
| | | 5 Others | N/A | N/A | N/A | > 0.05 |
| **GAAGGAGAAGAAGGAGAAGA** | 908 | **Protein Superfamily** | | | | |
| | | IPR017946 | PLC-like_Pdiesterase_TIM-brl | 6 | 4 | 1.94E-02 |
| | | 18 Others | N/A | N/A | N/A | > 0.05 |
| | | **Protein Family** | | | | |
| | | IPR045056 | Nop56/Nop58 | 142 | 141 | 7.90E-99 |
| | | IPR004802 | tRNA_PsdUridine_synth_B_fam | 82 | 81 | 1.99E-55 |
| | | IPR051270 | Tyrosine-tRNA_ligase_regulator | 27 | 27 | 2.81E-19 |

| | | | | | | |
|---|---|---|---|---|---|---|
| | | IPR024166 | rRNA_assembly_KRR1 | 22 | 22 | 8.04E-16 |
| | | IPR050656 | PINX1 | 37 | 29 | 7.81E-14 |
| | | IPR013240 | DNA-dir_RNA_pol1_su_RPA34 | 9 | 9 | 7.04E-07 |
| | | IPR053263 | Euk_RPA34_RNAP_subunit | 8 | 8 | 3.41E-06 |
| | | IPR026714 | SMAP | 13 | 8 | 1.60E-03 |
| | | IPR037804 | SGF73 | 3 | 3 | 8.97E-03 |
| | | IPR039328 | WDR89 | 3 | 3 | 8.97E-03 |
| | | IPR050420 | Calpain | 3 | 3 | 8.97E-03 |
| | | IPR001192 | PI-PLC_fam | 6 | 4 | 1.94E-02 |
| | | IPR039732 | Hub1/Ubl5 | 2 | 2 | 4.32E-02 |
| | | IPR040059 | PUM3 | 2 | 2 | 4.32E-02 |
| | | IPR007823 | RRP8 | 2 | 2 | 4.32E-02 |
| | | IPR050786 | EFG1_rRNA-proc | 2 | 2 | 4.32E-02 |
| | | 53 Others | N/A | N/A | N/A | > 0.05 |
| | | **Domain** | | | | |
| | | IPR000909 | PLipase_C_PInositol-sp_X_dom | 6 | 4 | 1.94E-02 |
| | | 12 Others | N/A | N/A | N/A | > 0.05 |
| | | **Repeats** | | | | |
| | | IPR008160 | Collagen | 2 | 1 | 3.73E-01 |
| | | **Other Features** | | | | |
| | | mobidb-lite | disorder_prediction | 3393 | 862 | 9.54E-56 |
| | | Coil | Coil | 2135 | 634 | 1.40E-46 |
| | | PTHR11467 | HISTONE H1 | 10 | 10 | 1.45E-07 |
| | | PTHR23236 | EUKARYOTIC TRANSLATION INITIATION FACTOR 4B/4H | 52 | 25 | 9.28E-06 |
| | | PTHR33840 | DUF2235 DOMAIN-CONTAINING PROTEIN | 4 | 4 | 1.86E-03 |
| | | PTHR37014 | EXPRESSION LETHALITY PROTEIN HEL10, PUTATIVE (AFU_ORTHOLOGUE AFUA_1G06580)-RELATED | 4 | 4 | 1.86E-03 |
| | | cd08598 | PI-PLC1c_yeast | 6 | 4 | 1.94E-02 |
| | | PS50007 | PIPLC_X_DOMAIN | 7 | 4 | 3.80E-02 |
| | | PTHR45815 | PROTEIN DISULFIDE-ISOMERASE A6 | 2 | 2 | 4.32E-02 |
| | | 32 Others | N/A | N/A | N/A | > 0.05 |
| **GACGGCTTCATGATCCACTC** | 361 | **Protein Superfamily** | | | | |
| | | IPR036419 | Ribosomal_S3_C_sf | 361 | 361 | 0.00E+00 |
| | | **Protein Family** | | | | |
| | | IPR005703 | Ribosomal_uS3_euk/arc | 358 | 357 | 0.00E+00 |
| | | **Domain** | | | | |
| | | IPR001351 | Ribosomal_uS3_C | 360 | 359 | 0.00E+00 |
| | | **Conserved Site** | | | | |
| | | IPR018280 | Ribosomal_uS3_CS | 351 | 351 | 0.00E+00 |
| | | **Other Features** | | | | |
| | | PTHR11760 | 30S/40S RIBOSOMAL PROTEIN S3 | 359 | 359 | 0.00E+00 |
| | | CYTOPLASMIC_DOMAIN | Cytoplasmic domain | 504 | 1 | 1.00E+00 |
| **GAGGCTGGTATCTCCAAGGA** | 732 | **Protein Superfamily** | | | | |
| | | IPR027417 | P-loop_NTPase | 1013 | 730 | 0.00E+00 |
| | | **Protein Family** | | | | |
| | | IPR050100 | TRAFAC_GTPase_members | 846 | 726 | 0.00E+00 |
| | | IPR004539 | Transl_elong_EF1A_euk/arc | 833 | 716 | 0.00E+00 |
| | | **Domain** | | | | |
| | | IPR000795 | T_Tr_GTP-bd_dom | 851 | 729 | 0.00E+00 |
| | | **Other Features** | | | | |
| | | cd01883 | EF1_alpha | 819 | 702 | 0.00E+00 |

| | | | | | | |
|---|---|---|---|---|---|---|
| | | ANF00225 | Spurious_ORF_225 | 2 | 2 | 2.81E-02 |
| | | 2 Others | N/A | N/A | N/A | > 0.05 |
| **TCCAACTTCATCAAGAAGGT** | 565 | **Protein Superfamily** | | | | |
| | | IPR027417 | P-loop_NTPase | 1013 | 565 | 0.00E+00 |
| | | **Protein Family** | | | | |
| | | IPR050100 | TRAFAC_GTPase_members | 846 | 563 | 0.00E+00 |
| | | IPR004539 | Transl_elong_EF1A_euk/arc | 833 | 561 | 0.00E+00 |
| | | **Domain** | | | | |
| | | IPR000795 | T_Tr_GTP-bd_dom | 851 | 564 | 0.00E+00 |
| | | **Other Features** | | | | |
| | | cd01883 | EF1_alpha | 819 | 549 | 0.00E+00 |
| | | 3 Others | N/A | N/A | N/A | > 0.05 |
| **TGTTGTTGTTGTTGTTGTGG** | 541 | **Protein Superfamily** | | | | |
| | | 35 Others | N/A | N/A | N/A | > 0.05 |
| | | **Protein Family** | | | | |
| | | IPR052099 | Regulatory_TF_Diverse | 5 | 5 | 2.87E-05 |
| | | IPR001289 | NFYA | 4 | 4 | 2.33E-04 |
| | | IPR050797 | Carb_Metab_Trans_Reg | 3 | 3 | 1.89E-03 |
| | | IPR017073 | HGS/VPS27 | 2 | 2 | 1.53E-02 |
| | | IPR027124 | Swc5/CFDP1/2 | 2 | 2 | 1.53E-02 |
| | | IPR040150 | Iwr1 | 2 | 2 | 1.53E-02 |
| | | IPR016966 | Thiamin_pyrophosphokinase_euk | 2 | 2 | 1.53E-02 |
| | | IPR006282 | Thi_PPkinase | 2 | 2 | 1.53E-02 |
| | | IPR051711 | Stress_Response_Reg | 2 | 2 | 1.53E-02 |
| | | IPR051059 | Fungal_TF_ZincFinger | 2 | 2 | 1.53E-02 |
| | | IPR027725 | HSF_fam | 2 | 2 | 1.53E-02 |
| | | IPR051079 | Sorting_Nexin_Autophagy | 2 | 2 | 1.53E-02 |
| | | IPR040309 | Naf1 | 2 | 2 | 1.53E-02 |
| | | IPR051374 | Ataxin-10/CTR86_families | 2 | 2 | 1.53E-02 |
| | | IPR039900 | Pat1-like | 2 | 2 | 1.53E-02 |
| | | IPR047767 | PSP1-like | 2 | 2 | 1.53E-02 |
| | | IPR004835 | Chitin_synth | 3 | 2 | 4.22E-02 |
| | | IPR053019 | GATA_zinc_finger | 3 | 2 | 4.22E-02 |
| | | 134 Others | N/A | N/A | N/A | > 0.05 |
| | | **Domain** | | | | |
| | | 25 Others | N/A | N/A | N/A | > 0.05 |
| | | **Repeats** | | | | |
| | | IPR025574 | Nucleoporin_FG_rpt | 4 | 2 | 7.75E-02 |
| | | **Other Features** | | | | |
| | | SSF81995 | beta-sandwich domain of Sec23/24 | 15 | 13 | 1.18E-10 |
| | | mobidb-lite | disorder_prediction | 3393 | 462 | 1.15E-06 |
| | | PTHR43329 | EPOXIDE HYDROLASE | 3 | 3 | 1.89E-03 |
| | | PTHR45735 | CLEAVAGE STIMULATION FACTOR SUBUNIT 2 | 3 | 3 | 1.89E-03 |
| | | SSF55961 | Bet v1-like | 2 | 2 | 1.53E-02 |
| | | PTHR13622 | THIAMIN PYROPHOSPHOKINASE | 2 | 2 | 1.53E-02 |
| | | PTHR47794 | VACUOLAR PROTEIN SORTING-ASSOCIATED PROTEIN 27 | 3 | 2 | 4.22E-02 |
| | | 80 Others | N/A | N/A | N/A | > 0.05 |
