## Supplementary Data 5 for "Kingdom-Wide CRISPR Guide Design with ALLEGRO"

**Phylogenetic tree highlighting the broader fungal species groups targeted by E1 sgRNAs, which were designed and validated for auxotrophic genes in *K. marxianus*, *K. phaffii*, and *Y. lipolytica*.**

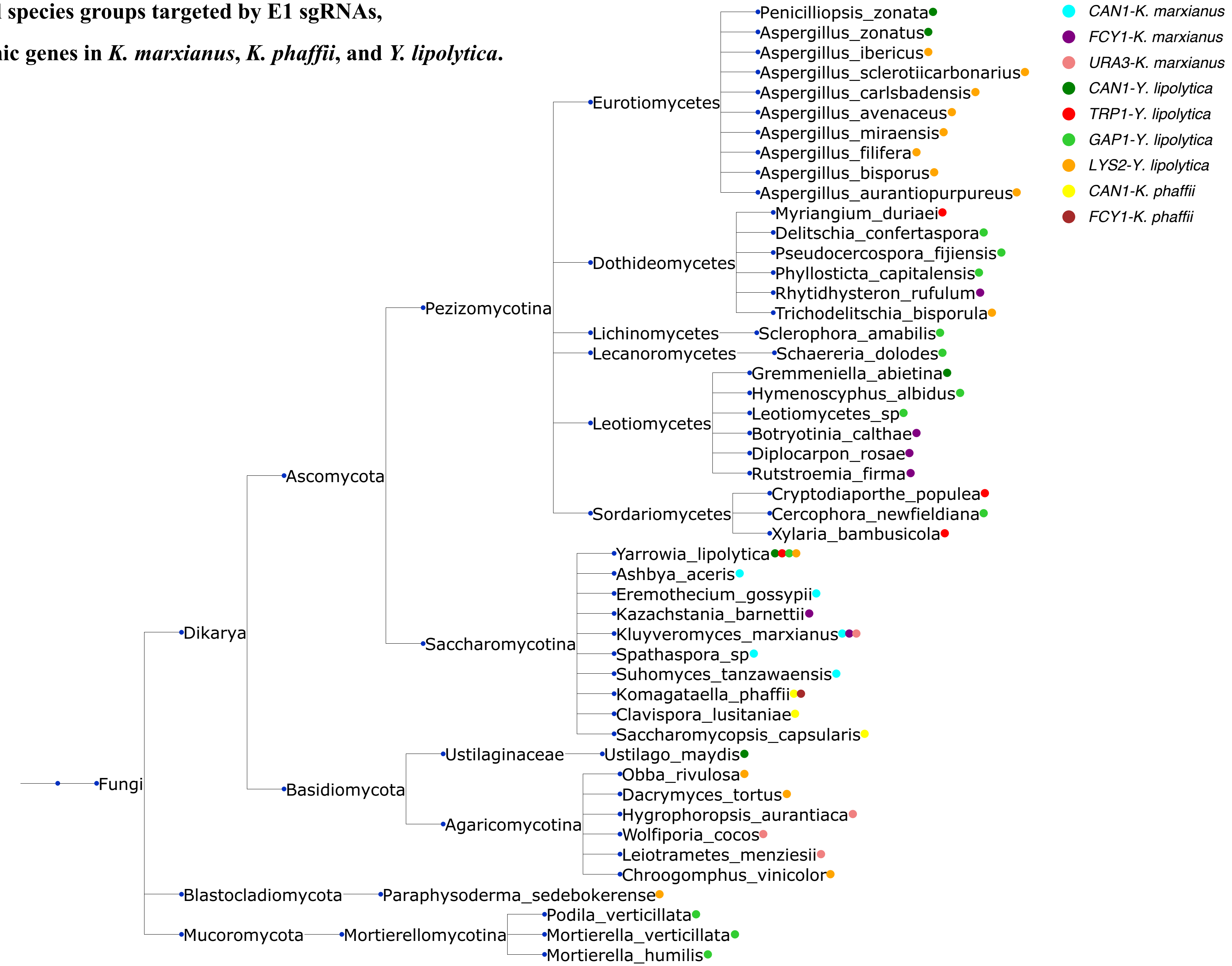
